# Diel fluctuations in *in-vivo* SnRK1 activity in Arabidopsis rosettes during light-dark cycles

**DOI:** 10.1101/2022.09.23.509272

**Authors:** Omri Avidan, Thiago A. Moraes, Virginie Mengin, Regina Feil, Filip Rolland, Mark Stitt, John E. Lunn

**Affiliations:** Max Planck Institute of Molecular Plant Physiology, Am Muehlenberg 1, 14476 Potsdam-Golm, Germany; Laboratory of Molecular Plant Biology, KU Leuven, Leuven, Belgium; KU Leuven Plant Institute (LPI), Leuven, Belgium

**Keywords:** *Arabidopsis thaliana*, SnRK1, trehalose 6-phosphate, glucose 6-phosphate, diel, circadian clock, starch

## Abstract

SUCROSE-NON-FERMENTING1 (SNF1)-RELATED KINASE1 (SnRK1) is a central hub in carbon and energy signalling in plants, and is orthologous with SNF1 in yeast and the AMP-ACTIVATED PROTEIN KINASE (AMPK) in animals. Previous studies of SnRK1 relied on *in-vitro* activity assays or on monitoring the expression of putative marker genes. Neither approach gives unambiguous information about *in-vivo* SnRK1 activity. We have monitored *in-vivo* SnRK1 activity using Arabidopsis (*Arabidopsis thaliana*) reporter lines that express a chimeric polypeptide with a SNF1/SnRK1/AMPK-specific phosphorylation site. We investigated responses during an equinoctial diel cycle, and after perturbing this cycle. As expected, *in vivo* SnRK1 activity rose towards the end of the night and rose even further when the night was extended. Unexpectedly, although sugars rose after dawn, SnRK1 activity did not decline until about 12 hours into the light period. The sucrose signal trehalose 6-phosphate (Tre6P) has been shown to inhibit SnRK1 *in vitro*. We introduced the SnRK1 reporter into lines that harboured an inducible *TREHALOSE-6-PHOSPHATE SYNTHASE* construct. Elevated Tre6P decreased *in-vivo* SnRK1 activity in the light period, but not at the end of the night. Reporter polypeptide phosphorylation was sometimes negatively correlated with Tre6P, but a stronger and more widespread negative correlation was observed with glucose 6-phosphate. We propose that SnRK1 operates within a network that controls carbon utilization and maintains diel sugar homeostasis, and that Tre6P, hexose phosphates and the circadian clock contribute to regulation of SnRK1 activity in a context-dependent manner, and SnRK1-signalling is further modulated by factors that act downstream of SnRK1.

**One sentence summary:** *In vivo* SnRK1 activity shows an unexpected diel response and a complex relationship with trehalose 6-phosphate and other possible metabolic regulators.

## Introduction

Plant SUCROSE-NON-FERMENTING1-RELATED PROTEIN KINASES 1 (SnRK1) belongs to a eukaryotic family of Ca^+2^-independent Ser/Thr protein kinases. SnRK1 is implicated in maintaining cellular homeostasis and energy balance alongside linking carbon (C) availability and growth, and has been extensively studied in recent years (Polge and Thomas, 2006; Jossier *et al*., 2009; Hulsmans *et al*., 2016; Baena-González *et al*., 2007; Paul *et al*., 2018; 2020; Crepin and Rolland, 2019; Baena-González and Lunn, 2020). These functions are broadly similar to those observed for its yeast and animal orthologues, SUCROSE-NON-FERMENTING1 (SNF1) and AMP-ACTIVATED PROTEIN KINASE (AMPK) but our understanding of the functions and regulation of SnRK1 in plants is much less complete.

SnRK1 is a heterotrimer consisting of a catalytic (α) subunit and two regulatory (β and βγ) subunits that modify complex stability, activity, interactions and cellular localization (Polge *et al*., 2008; Nietzsche *et al*., 2016; Wang *et al*., 2020). It is located in the nucleus and cytosol (Nietzsche *et al*., 2014; Blanco *et al*., 2019; Ramon *et al*., 2019) and has also been reported to be present in plastids (Ávila-Castañeda *et al*., 2014; Ruiz-Gayosso *et al*., 2018). According to current knowledge, the mode of regulation differs from that of SNF1 and AMPK. The regulatory properties of these kinases have been studied mainly by *in-vitro* assays, which monitor phosphorylation of peptides containing an evolutionarily conserved SNF1/SnRK1/AMPK-specific recognition sequence: [MLVFI].X.[RKH].X.X.**<u>S</u>**.X.X.X.[LFIMV] or the less favoured [MLVFI].[RKH].X.X.X.**<u>S</u>**.X.X.X.[LFIMV] (Dale *et al*., 1995). Such studies have demonstrated that animal AMPK, and to a certain degree also yeast SNF1, are directly regulated by AMP and/or ADP (Hardie and Carling, 1997; Hardie *et al*., 1998; Mayer *et al*., 2011). In mammals, these adenylates inhibit T-loop dephosphorylation, thereby activating the catalytic (α) subunit in low energy conditions and/or in response to low C availability (García-Salcedo *et al*., 2014; Crozet *et al*., 2014). In contrast, plant SnRK1 is not activated by adenylates in *in-vitro* assays. Instead, it is inhibited *in vitro* by sugar phosphates, such as trehalose 6-P (Tre6P), glucose 6-phosphate (Glc6P) and glucose 1-phosphate (Glc1P) (Toroser *et al*., 2000; Lu *et al*., 2007; Zhang *et al*., 2009; Emanuelle *et al*., 2015; Broeckx *et al*., 2016; Zhai *et al*., 2017; 2018; Ruiz-Gayosso *et al*., 2018). Inhibition by sugar phosphates could potentially link SnRK1 activity to C availability rather than energy status (Hara *et al*., 2013; Nunes *et al*., 2013; Herzig and Shaw, 2017). These *in-vitro* studies also indicated that inhibition by Tre6P requires an unknown proteinaceous factor that is present in growing tissues but not in mature source leaves (Toroser *et al*., 2000; Zhang *et al*., 2009; Emanuelle *et al*., 2015). Subsequent studies have shown that Tre6P can bind directly to the SnRK1 catalytic α subunit, thereby interfering with binding of the SnRK1-activating kinases GRIK1/2, leading to inhibition of SnRK1 activity (Glab *et al*., 2017; Zhai *et al*., 2018; Hwang et al., 2019). However, GRIK1/2 might not be the only mechanism by which Tre6P inhibits SnRK1 activity. As GRIK1/2 are widely expressed in source leaves, Zhai et al. (2018) concluded that they were unlikely to be the unknown proteinaceous factor from young tissues that was referred to by Zhang et al. (2009).

Glc6P and Glc1P are intermediates in glycolysis, gluconeogenesis and in the synthesis and degradation of major carbohydrate reserves like starch and sucrose. Tre6P is an intermediate in the pathway of trehalose biosynthesis. It is synthesized from UDP-Glc and Glc6P by TREHALOSE-6-PHOSPHATE SYNTHASE (TPS) and is dephosphorylated to trehalose by TREHALOSE-6-PHOSPHATE PHOSPHATASE (TPP) (Cabib and Leloir, 1958). Trehalose is a major C storage metabolite, osmolyte and stress protectant in fungi, but has been largely replaced by sucrose in vascular plants, allowing neo-functionalization of the trehalose biosynthesis pathway (Goddijn and van Dun, 1999; Lunn *et al*., 2014; Lunn, 2016). In addition to catalytically active class I TPS proteins), plants possess a large family of catalytically inactive class II TPS proteins (Leyman *et al*., 2001; Harthill *et al*., 2006; Lunn, 2007; Vandesteene *et al*., 2010; Lunn et al., 2014; Delorge *et al*., 2015) as well as a large family of TPP proteins (Leyman *et al*., 2001; Vandesteene *et al*., 2012; Kretzschmar *et al*., 2015). Tre6P is considered to serve as a signal of sucrose availability (Lunn *et al*., 2006), acting to regulate C allocation between sucrose and other photosynthesis products in source leaves, and to link sucrose supply and growth in sink organs (Schluepmann *et al*., 2003; Martins *et al*., 2013; Wahl *et al*., 2013; Figueroa *et al*., 2016; Paul *et al*., 2018; 2020; Fichtner and Lunn, 2021).

The possible contribution of Glc6P and Glc1P to regulation of SnRK1 activity *in planta* has not been addressed in detail. Interactions between Tre6P and SnRK1 have been more extensively studied, mainly by investigating changes in SnRK1 marker transcript abundance (Zhang *et al*., 2009; 2015; Henry *et al*., 2015; Bledsoe *et al*., 2017) and post-translational regulation of target enzymes Baena-González and Sheen, 2008; Figueroa *et al*., 2016). The former builds on the work of Baena-González *et al*. (2007) who, using a protoplast overexpression system, identified over 1000 genes as downstream targets of SnRK1, such as *DIN6*, *DIN1*, *BCAT2* and *EXP10*, whose expression is regulated by SnRK1-mediated phosphorylation of upstream transcription factors, such as bZIP11 (Ma et al., 2011) and bZIP63 (Mair et al., 2016). Their expression has been widely used as a sensitive readout of SnRK1 activity, either via promoter activation assays (Rodrigues *et al*., 2013; Ramon *et al*., 2019) or by analyses of transcript abundance. Abundance of SnRK1-induced transcripts is typically low in a diel cycle but rises strongly when plants are starved by transfer to continuous darkness (Baena-González *et al*., 2007; Baena-González and Sheen, 2008; Usadel *et al*., 2008; Flis *et al*., 2016) as expected if SnRK1 is activated in C starvation conditions (Mair *et al*., 2015; Pedrotti *et al*., 2018; Ramon *et al*., 2019). Their abundance often changes reciprocally to Tre6P levels in starvation treatments and other perturbations in several species and tissues (Martínez-Barajas *et al*., 2011; Wingler *et al*., 2012; Nunes *et al*., 2013; Henry *et al*., 2015; Nuccio *et al*., 2015; Baena-González and Lunn, 2020; Peixoto *et al*., 2021) providing correlative evidence for the idea that Tre6P inhibits SnRK1 *in vivo*. Application of permeable Tre6P analogues to Arabidopsis led to changes in transcript abundance for a subset of these SnRK1 downstream target genes that were consistent with Tre6P acting to inhibit SnRK1 (Griffiths *et al*., 2016). Further support for partial convergence of the SnRK1 and Tre6P signalling pathways was provided by the identification of several class II TPSs as transcriptional and post-translational SnRK1 targets (Glinski and Weckwerth, 2005; Harthill *et al*., 2006; Cho *et al*., 2016; Nukarinen *et al*., 2016). However, the functional significance of this observation remains unclear because the precise role of the TPS class II proteins has not yet been defined (Schluepmann and Paul, 2009; Ponnu *et al*., 2011; Yang *et al*., 2012; Lunn et al. 2014; Delorge *et al*., 2015; Figueroa and Lunn, 2016). More recently, it was demonstrated that mutant lines with increased or reduced expression of SnRK1 exhibit an altered relationship between sucrose and Tre6P in diel cycles, as well as modified anaplerotic flux of carbon to the tricarboxylic acid cycle that is reminiscent of responses to Tre6P (Peixoto *et al*., 2021). Furthermore, mutations in the *SnRK1α1* and *SnRK1βγ* genes were shown to suppress developmental defects in the Arabidopsis *tps1* mutant (Zacharaki *et al*., 2022).

Taken together, these results point to SnRK1 and Tre6P signalling being closely intertwined. They also indicate that SnRK1 signalling is involved not only in starvation responses, but also in the coordination of metabolism in benign conditions where plants are not C-starved or facing a large C excess (see also Baena-González and Lunn, 2020). However, many open questions remain; for example, negative correlations between Tre6P levels and the transcript abundance of SnRK1-induced genes are seen in mature tissues even though Tre6P does not inhibit SnRK1 activity in *in-vitro* assays with material from mature tissues (see above and Baena-González and Lunn, 2020). Moreover, there is an inconsistent correlation between Tre6P levels and transcript abundance of SnRK1 target genes in *in-vitro* cultured excised maize kernels (Bledsoe *et al*., 2017, see also Baena-González and Lunn, 2020). It remains possible that Tre6P also acts on signalling downstream of SnRK1, as is the case in yeast (Deroover *et al*., 2016), resulting in changes in downstream transcript abundance that are not directly due to inhibition of SnRK1 itself. Interpretation is further complicated by potential interactions between SnRK1 and TARGET OF RAPAMYCIN (TOR) signalling (Lastdrager *et al*., 2014; Margalha and Confraria, 2019; Ryabova *et al*., 2019).

The two main approaches employed in plants to study SnRK1 regulation have shortcomings which have not been addressed to date. The *in-vitro* assay is direct and highly specific, but the use of crude extracts or purified proteins may bring about other complications due to loss of the *in-vivo* context, such as artificial interactions or potential loss of regulatory elements that otherwise would have an effect on activity (Baena-González and Lunn, 2020). Monitoring changes in expression of SnRK1 marker genes is sensitive, but gene expression is an indirect readout that may be prone to secondary effects that are unrelated to SnRK1 activity. Indeed, there is often poor agreement between SnRK1 activity in the *in-vitro* phosphorylation assay and changes in SnRK1 target gene expression (Debast *et al*., 2011; Martínez-Barajas *et al*., 2011; Baena-González and Lunn, 2020).

Working in yeast, Deroover *et al*. (2016) developed a specific *in-vivo* SNF1 reporter that harbours an SNF1/SnRK1/AMPK-specific recognition sequence-containing peptide, based on rat ACETYL COA CARBOXYLASE 1 (ACC1), an established AMPK target (Dale *et al*., 1995). The phosphorylation status of this peptide can be monitored by extraction and immunoblot analysis with a validated commercial phospho-specific antibody, providing a direct measure of *in-vivo* SNF1 activity. Subsequently, modified ACC1 reporters were used in Arabidopsis to study the regulation of SnRK1 activity by nitrogen status (Sanagi *et al*., 2021) and in other responses (Muralidhara *et al*., 2021;cHenninger *et al*., 2022; Belda-Palazon *et al*., 2022).

We have used Arabidopsis ACC1 reporter lines to investigate *in-vivo* SnRK1 activity during a diel cycle in benign conditions, when the C supply is only slightly restricting growth, and after changing the light regime to modify the C supply. We have also introduced the reporter construct into Arabidopsis lines in which Tre6P levels can be increased in an inducible manner (Martins *et al*., 2013; Figueroa *et al*., 2016). We show that SnRK1 activity does not change reciprocally to overall C availability, with a peak at dawn and a minimum at dusk. Instead, SnRK1 activity appears to be maintained in the light period, declines after darkening and rises towards the end of the night. Furthermore, SnRK1 activity *in vivo* is not strongly or consistently inhibited by an induced increase in Tre6P. Multiple regression analyses of metabolite levels indicate that the link between *in-vivo* SnRK1 activity and the level of Tre6P is context-dependent and point to SnRK1 activity being regulated *in vivo* by additional mechanisms, including inhibition by Glc6P.

## Results

### Reporter constructs to quantify in vivo SnRK1 activity

To obtain a direct *in-vivo* readout of SnRK1 activity in Arabidopsis we employed transgenic SnRK1-reporter lines expressing a peptide derived from the Ser79 phosphorylation site of ACC1 fused to GREEN FLUORESCENT PROTEIN (GFP), which was used to normalize for the level of expression of the reporter polypeptide (see Supplementary Figure S1 for full description) and to monitor its localization (see below). The level of phosphorylation of the ACC1 peptide (pACC) was assessed using a specific anti-phospho-site antibody, and an anti-GFP antibody was used to normalize for expression level and gel loading. To increase statistical and biological robustness, we used three-four biological replicates per time point. An additional control sample (extended night) was loaded in all gels, and used to correct for between-gel variation. We took the normalized pACC/GFP signal ratio as a readout of *in-vivo* SnRK1 activity at a given time point (for details see Methods and Supplementary Figure S1).

Two transgenic lines were used: (i) expressing a nuclear reporter (NUC), that contains the SV40 nuclear location signal (NLS) and (ii) expressing a general reporter (GEN), in which GFP is observed in both the nucleus and the cytoplasm (Supplementary Figure S2).

### Validation of the reporter construct in Arabidopsis rosettes

The specificity of the ACC1 reporter peptide was previously tested in yeast by incubating WT and *snf1* mutant cells with fermentable glycerol and ethanol to induce SNF1 or with glucose to inhibit SNF1 activity (Deroover *et al*., 2016). The peptide was phosphorylated under fermenting conditions but not following glucose addition in the wild type background, while no phosphorylation was observed in the *snf1* background, suggesting that the reporter is indeed SNF1-specific in yeast. Sanagi *et al*. (2021) validated an ACC1-based SnRK1 reporter in Arabidopsis mesophyll protoplasts, by showing that phosphorylation is increased by transient expression of wild-type SnRK1α1 but not by expression of a mutated form, SnRK1α1^K48A^, which is defective in the SnRK1 ATP-binding pocket (K48M). They also showed that there is less phosphorylation in protoplasts prepared from inducible *snrk1α* knockdown plants than in protoplasts prepared from wild-type plants.

We took two approaches to further validate the reporters for use in whole Arabidopsis rosettes. In the first approach, we subjected GEN and NUC lines to 3-h of extended darkness, starting at the end of the night (Supplementary Figure S3A; Supplementary Dataset S1 – Exp.1). Extended darkness is known to activate SnRK1 (see Introduction). As expected, phosphorylation of both reporters increased in extended darkness, with phosphorylation being significantly higher at ZT25 (zeitgeber time, i.e. 25 h after the previous dawn and into the extended night) and rising further at ZT27 in NUC, and being significantly higher by ZT27 in GEN. In the second approach, the NUC reporter was crossed to a 35:SnRK1α1 line that has elevated SnRK1 activity and to the 35S:SnRK1α1^K48M^ line expressing a catalytically inactive mutant protein (Jossier *et al*., 2009; Crozet *et al*., 2014; Cho *et al*., 2016) (Supplementary Figure S3B; Supplementary Dataset S1 – Exp.2). The crossed lines and a NUC control were grown in equinoctial growth conditions for 22 days after sowing (DAS), and then harvested at ZT0, 8, 16, 24 and 28 (i.e. 4-h of extended darkness). We confirmed expression of SnRK1 proteins in these lines by immunoblotting with anti-SnRK1α1 antibody (Supplementary Figure S3C). Phosphorylation of the NUC polypeptide was consistently higher in NUC х 35S:SnRK1α1 than in NUC х 35S:SnRK1^K48M^ (significant at ZT0, 8, 16, 24 and 28). Compared to the NUC control, reporter phosphorylation in NUC х 35S:SnRK1^K48M^ was significantly lower at ZT0, 8, 24 and 28 (with SnRK1^K48M^ apparently functioning as a dominant negative mutant protein; Cho et al., 2016), and reporter phosphorylation in NUC х 35S:SnRK1α1 was essentially control-like at ZT0, 8, 24 and 28 and significantly higher at ZT16. This experiment also confirmed the rise in reporter phosphorylation in the extended night, further validating the reporter.

### Diel changes in SnRK1 activity in equinoctial growth conditions

The diel response of the NUC control in Supplementary Figure S3B revealed a somewhat unexpected pattern, with the signal being quite high throughout the light period, decreasing rapidly at the beginning of night, and then rising in the last part of the night. This was unexpected as photosynthesis provides a higher C supply in the light, which is expected to inhibit SnRK1. We investigated if this unexpected diel response was robust and reproducible. Independent experiment with equinoctial light-dark regimes (three with NUC, two with GEN) yielded a robust and reproducible diel response with SnRK1 activity at dawn being maintained during the light period, followed by a decline after darkening to a minimum at about ZT16 and an increase in the last part of the night (Figure 1A). The NUC and GEN lines exhibited similar responses.

**Figure 1:**
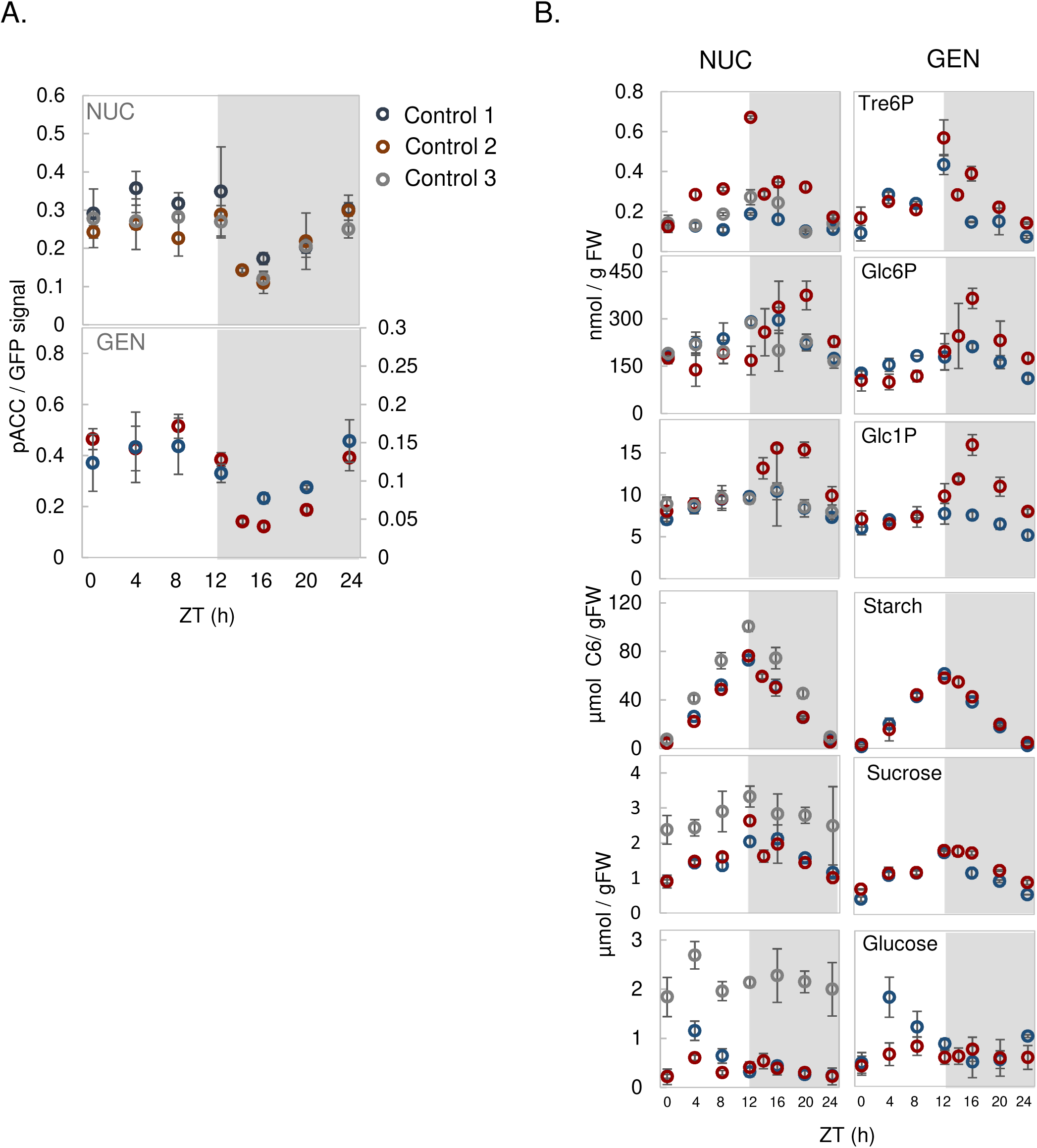
Reproducible diel response in phosphorylation of SnRK1 reporters under equinoctial growth conditions. SnRK1 reporter lines were grown in a 12-h photoperiod (irradiance 160 µmol m^-2^ s^-1^) for 22 days as controls in three independent experiments, in which additional plants grown in parallel were subjected to different treatments (see Figs 2-4). Whole rosettes were harvested at 4-h intervals through the diel cycle. (A) Phosphorylation of the NUC/GEN polypeptide was quantified by immunoblotting as described in Methods and in Supplementary Figure S1. (B) Selected metabolites. Data are from the control samples from the following experiments: (i) Control 1: Supplementary Dataset S1 – Exp.3 (blue symbols; NUC and GEN lines); (ii) Control 2: Supplementary Dataset S1 – Exp.4 and Supplementary Figure S4 (red symbols; NUC and GEN lines); (iii) Control 3: Supplementary Dataset S1 – Exp.2 and Supplementary Figure S3b (grey symbols; NUC line only). Data are shown as mean ± S.D. (n = 3-4 biological replicates, each containing 3-5 pooled rosettes). Statistical analyses are provided for each time series in Figs 2, 3 and 4, respectively.

We explored diel changes in metabolic status and in potential metabolite effectors of SnRK1 (Figure 1B) in the same plant material. Obvious candidates include sugars (glucose, sucrose) and starch as indicators of C status, and sugar phosphates, such as Tre6P, Glc6P and Glc1P, all of which were suggested previously to be directly or indirectly involved in the regulation of SnRK1 (Toroser *et al*., 2000; Lu *et al*., 2007; Zhang *et al*., 2009; Emanuelle *et al*., 2015; Broeckx *et al*., 2016; Hulsmans *et al*., 2016; Zhai *et al*., 2017; 2018; Ruiz-Gayosso *et al*., 2018). The diel responses of metabolites were reproducible and resembled previous reports (Lunn *et al*., 2006; Sulpice *et al*., 2014; Mengin *et al*., 2017; Flis *et al*., 2019; Moraes *et al*., 2019). In brief, starch accumulated in a near-linear manner during the light period and was mobilized in a near-linear manner at night, sucrose gradually increased in the light to a peak at ZT12 (ED) and gradually decreased during the night, and glucose levels showed a 1.5- to 3-fold transient increase at ZT4 but otherwise remained relatively low throughout the diel cycle (see also Mengin *et al*., 2017). Tre6P changed in parallel with sucrose, as expected from previous studies (Martins *et al*., 2013; Figueroa *et al*., 2016; Peixoto *et al*., 2021). Glc6P and Glc1P levels remained fairly stable throughout the day, rose at the beginning of the night until ZT16 and then decreased.

These experiments did not reveal pronounced reciprocal changes in Tre6P levels and SnRK1 activity during the diel cycle, except late in the night (between ZT16-24) when Tre6P levels decreased and phosphorylation of both NUC and GEN polypeptides increased. In addition, no reciprocal changes were observed between NUC or GEN phosphorylation and sugar levels, except for sucrose in the last part of the night. However, Glc6P and Glc1P did show a reciprocal relationship with SnRK1 activity through the night; Glc6P and Glc1P initially increased after dusk (from ZT12-16), and then decreased (from ZT16-24), while phosphorylation of NUC and GEN decreased during the first part of the night and then increased (see later for more analysis).

### Perturbations of the light regime

To further explore how SnRK1 is regulated by changes in metabolites, by light and/or by endogenous (e.g. circadian clock) rhythms, we next investigated NUC and GEN phosphorylation during perturbations of the light regime, including standard equinoctial light treatments as a control.

The first treatment was based on a perturbation described in Moraes *et al*. (2019). Plants were grown in equinoctial conditions under 160 µmol m^-2^ s^-1^ irradiance for 19 days, and then transferred to 60 µmol m^-2^ s^-1^ for one light period (treatment light period), darkened at ZT12 at the same time as in the initial growth conditions (treatment night) and then returned to 160 µmol m^-2^ s^-1^ on the next day (recovery light period). Control plants were left in the original growth conditions (Figure 2). The response of starch and sugars closely resembled that in Moraes *et al*. (2019). Compared to control plants, a single low-irradiance day led to slower starch accumulation and a decrease in the levels of sucrose and glucose during the treatment light period, and slower starch mobilization and lower sugar levels in the following night (Figure 2A; Supplementary Dataset S1 – Exp.3). During the recovery light period, glucose was higher and starch accumulation was faster than in the light period in control plants, indicating that growth was restricted (see also Gibon *et al*., 2004; Sulpice *et al*., 2014b; Mengin *et al*., 2017).

**Figure 2:**
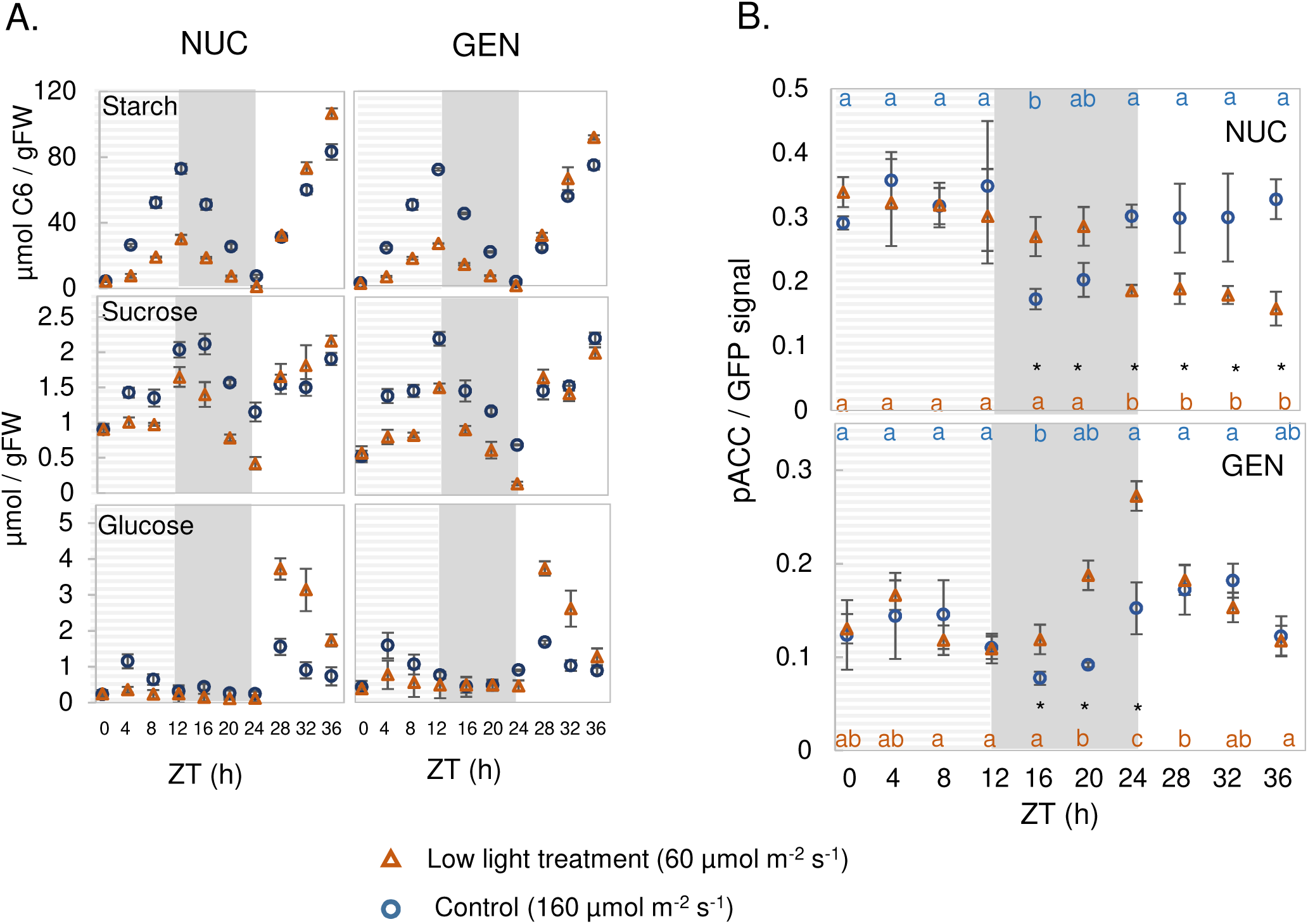
SnRK1 activity in Arabidopsis plants transferred to low irradiance for one light period. NUC and GEN lines were grown in a 12-h photoperiod at 160 µmol m^-2^ s^-1^ irradiance for 19 days. At 20 DAS, starting at dawn (ZT0), irradiance was reduced to 60 µmol m s for the following 12 h (treatment light period), plants were darkened at ZT12 (treatment night), and at ZT24 were re-illuminated at an irradiance of 160 µmol m^-2^ s^-1^ (recovery day), as described in Moraes *et al*. (2019). Samples from both genotypes and from the corresponding control treatment (plants left in the original growth light regime) were harvested at 4-h intervals from ZT0-ZT36 (with respect to dawn on the treatment day). (A) Soluble sugars and starch. (B) Phosphorylation of NUC (top) and GEN (bottom) SnRK1 reporter polypeptides. The dashed grey background denotes the low-light treatment day. Results are shown as mean ± S.D. (n=3-4 biological replicates). Statistical analysis: letters indicate significant (*P*<0.05) changes between different times for a given genotype and asterisks indicate significant (*P*<0.05) differences between lines at a given time point, according to one-way ANOVA and pairwise multiple comparison post-testing using the Holm-Sidak method. The original and additional data are available in Supplementary Dataset S1 – Exp.3. The control time-series is also shown in Figure 1 (blue symbols).

Phosphorylation of NUC and GEN polypeptides (Figure 2B) showed a combination of expected and unexpected features, which were partially shared between the NUC and GEN lines but also showed some differences. Rather surprisingly, a sudden decrease in irradiance had no immediate impact on NUC or GEN phosphorylation in the treatment light period. However, it did have significant effects in the following night and during the recovery day. For NUC, there was no initial drop in phosphorylation after darkening, with the result that phosphorylation was significantly higher than the control (where it decreased in the first part of the night). However, after ZT20, there was a sharp decrease in NUC phosphorylation and this persisted during the recovery light period. For GEN, phosphorylation increased significantly compared to the control by ZT16, and increased even further as the night progressed, followed by a decrease to control levels in the recovery light period.

It is rather surprising that, after a shift to low light, the decrease of sugar levels was not accompanied by an increase in NUC or GEN phosphorylation in the light period. This indicates that at this time in the diel cycle, nuclear and extra-nuclear SnRK1 activity are not responding to current C availability as reflected by overall sugar levels. At night, NUC and GEN phosphorylation were higher in plants exposed to low irradiance during the preceding light period than in the control, reflecting the lower C availability, but this response was much more marked for GEN than NUC. Indeed, NUC signal fell at the end of the night, even though sugars remained low. In the recovery day, when sugars were higher than in the control, NUC phosphorylation was lower than the control as would be expected if it were responding to current C availability. However, GEN phosphorylation was high, indicating that extra-nuclear SnRK1 activity was not responding to current C availability. Indeed, considering that GEN is a composite of nuclear and of cytoplasmic activity, it is likely that cytoplasmic SnRK1 activity was actually increased since NUC signal activity was decreased at this time (see above).

The second treatment was based on an experiment reported by Fernandez *et al*. (2017), in which plants were grown in equinoctial conditions with 160 µmol m^-2^ s^-1^ irradiance for 19 days and then transferred to continuous lower irradiance. The original study of Fernandez *et al*. (2017) investigated starch degradation in the light, and the low irradiance was used to avoid over-accumulation of starch in continuous light. The rationale for the current experiment was to test whether the diel changes in phosphorylation activity of SnRK1 are driven by the external light regime or by endogenous cues.

We grew plants in standard equinoctial conditions at 160 µmol m^-2^ s^-1^, then transferred them, at dawn on day 20 after sowing, to continuous light with an irradiance of 90 µmol m^-2^ s^-1^, and sampled them at 4-h intervals for 32 h (Figure 3). Control plants were left in the original growth conditions. After transfer to continuous low light, starch accumulated for about 12 h, although more slowly than in control plants, and starch levels then plateaued or even declined slightly in the subjective night from ZT12 onwards (Figure 3A; Supplementary Dataset S1 – Exp.4). Sugar levels were lower than in the control in the subjective day period (ZT0-12) but rose to reach higher levels from ZT16 onwards. These changes of starch and sucrose reflect a gradual but progressive increase in the rate of starch mobilization from about ZT10 onwards (see Fernandez *et al*., 2017 for details). Compared to the control, Glc6P was only marginally and non-significantly lower in the subjective day, and was consistently lower in the subjective night (Supplementary Figure S4). Once again, we observed that lower sugar levels during the first 12 h of the 24-h cycle were not accompanied by a significant increase in phosphorylation of NUC or GEN, which essentially exhibited a control-like response until ZT12 (Figure 3; Supplementary Dataset S1 – Exp.4). In both lines, SnRK1 activity in continuous low light was higher in the subjective night than in the control during the night. Nevertheless, the decline in SnRK1 activity that is observed after darkening control plants at ZT12 was also observed under low continuous light, although in a weakened form. NUC phosphorylation showed a gradual significant decrease (relative to ZT12) from ZT14 to ZT24, while GEN phosphorylation started to decrease by ZT12 and declined until ZT16 (Figure 3B). In the next subjective day, phosphorylation levels of both NUC and GEN resembled those in the control.

**Figure 3:**
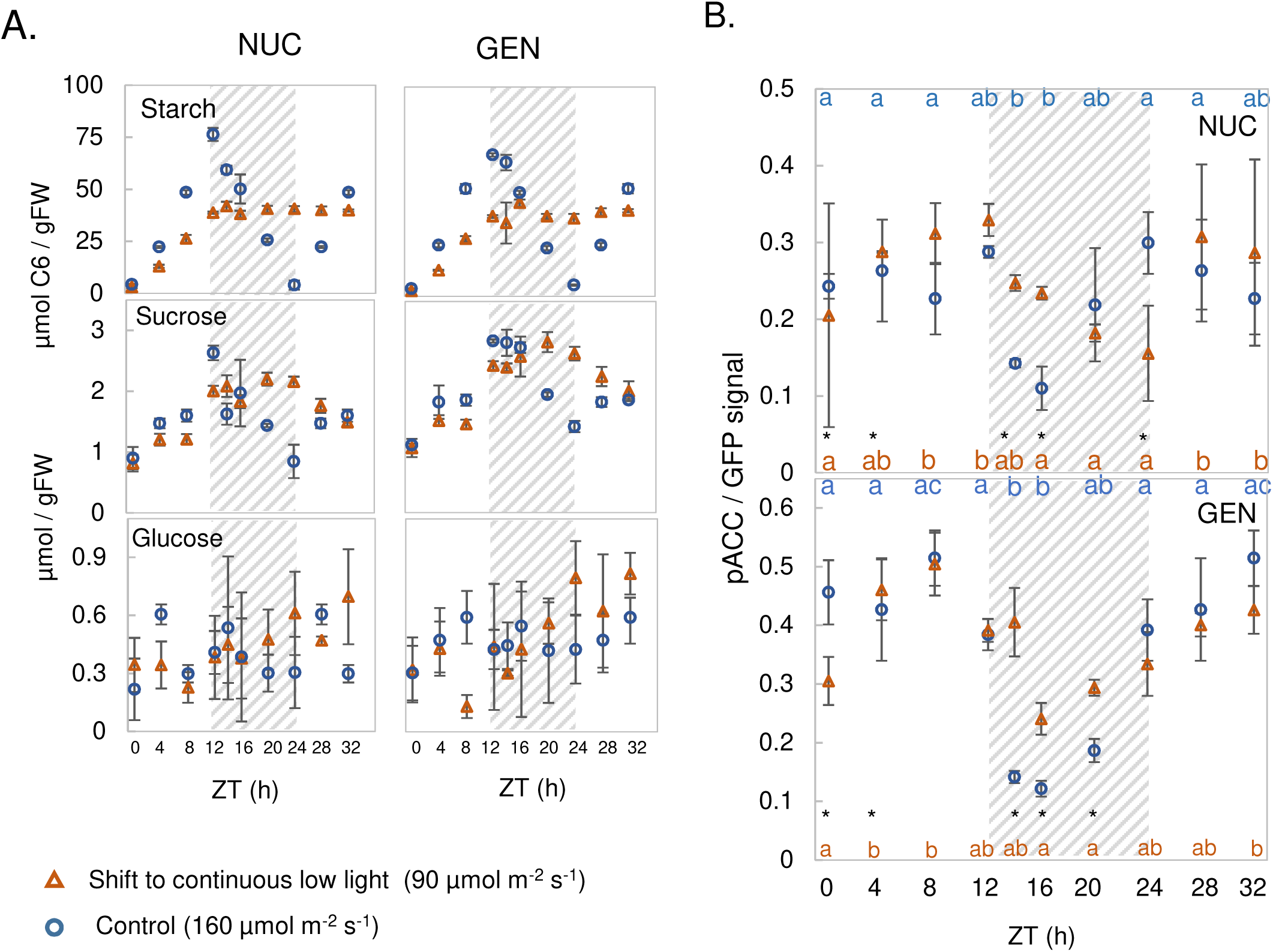
SnRK1 activity in Arabidopsis plants transferred to continuous low light. NUC and GEN lines were grown in a 12-h photoperiod at 160 µmol m^-2^ s^-1^ irradiance for 19 days. At 20 DAS, plants were shifted at dawn (ZT0) to continuous low light (90 µmol m^-2^ s^-1^) for 32 h, while corresponding controls were kept in the original growth conditions (12-h photoperiod, 160 µmol m^-2^ s^-1^). (A) Diel changes of sugars and starch. (B) Phosphorylation of NUC (top) and GEN (bottom) SnRK1 reporter polypeptides. The dashed grey background represents the subjective night. Results are shown as mean ± S.D. (n = 3 biological replicates). Statistical analysis: letters indicate significant (*P*<0.05) changes between different times for a given treatment and asterisks represent significant (*P*<0.05) differences between the low light treatment and the control at a given time point according to one-way ANOVA with pairwise multiple comparison post-testing using the Holm-Sidak method. The original and additional data are available in Supplementary Dataset S1 – Exp.4. The control time-series is also shown in Figure 1 (red symbols).

These observations confirmed that a fall in C availability due to decreased irradiance does not lead to increased SnRK1 activity in the first part of the 24-h cycle. They also point to an endogenous component contributing to the decline in SnRK1 activity from about ZT12 onwards. Furthermore, we noted a reciprocal relationship between diel changes in Glc6P and SnRK1 activity, particularly with NUC phosphorylation.

### Response in a T28 cycle

The Arabidopsis circadian clock is mainly entrained by light at dawn and imposes an endogenous 24-h rhythmicity even when the external cycle deviates from 24 h (Millar *et al*., 1995; Locke *et al*., 2005; Graf *et al*., 2010). Growing plants in a light-dark cycle that deviates from 24 h is therefore a good way to search for oscillatory responses that are driven directly or indirectly by the clock.

We grew plants in a 14-h light / 14-h dark (T28) or a standard 12-h light / 12-h dark (T24) cycle for 19 days with 160 µmol m^-2^ s^-1^ irradiance and harvested them at 4-h intervals on day 20 after sowing, including a 4-h or 8-h period of extended darkness for the T28 and T24 control, respectively (Figure 4). As previously seen (Graf *et al*., 2010), in a T28 cycle starch is exhausted at about ZT24, resulting in transient C starvation and low sucrose levels from ZT24 to ZT28 (i.e., the four hours preceding the next dawn) (Figure 4A; Supplementary Dataset S1 – Exp.5) as well as lower Glc6P levels through the night (Supplementary Figure S5A). Correspondingly, phosphorylation of NUC and GEN at ZT0 (dawn) was almost 6- and 3-fold higher in the T28 cycle than the T24 cycle, comparable to the increase observed after extending the night in a T24 cycle (Figure 4B; Supplementary Dataset S1 – Exp.5, see also Supplementary Fig. S3A). After illumination, NUC and GEN phosphorylation decreased strongly in a T28 cycle, falling to levels that were lower than in a T24 cycle for most of the remaining light-dark cycle (significant for both NUC and GEN at ZT8, ZT14 and ZT24). Importantly, NUC and GEN phosphorylation decreased between ZT12 and ZT14 in a similar manner in both the T28 and T24 cycle plants, even though the former were still in the light while the latter were already in the dark. Together with the response in continuous light (see above), these observations point to a component with 24-h rhythmicity contributing to the regulation of SnRK1 activity.

**Figure 4:**
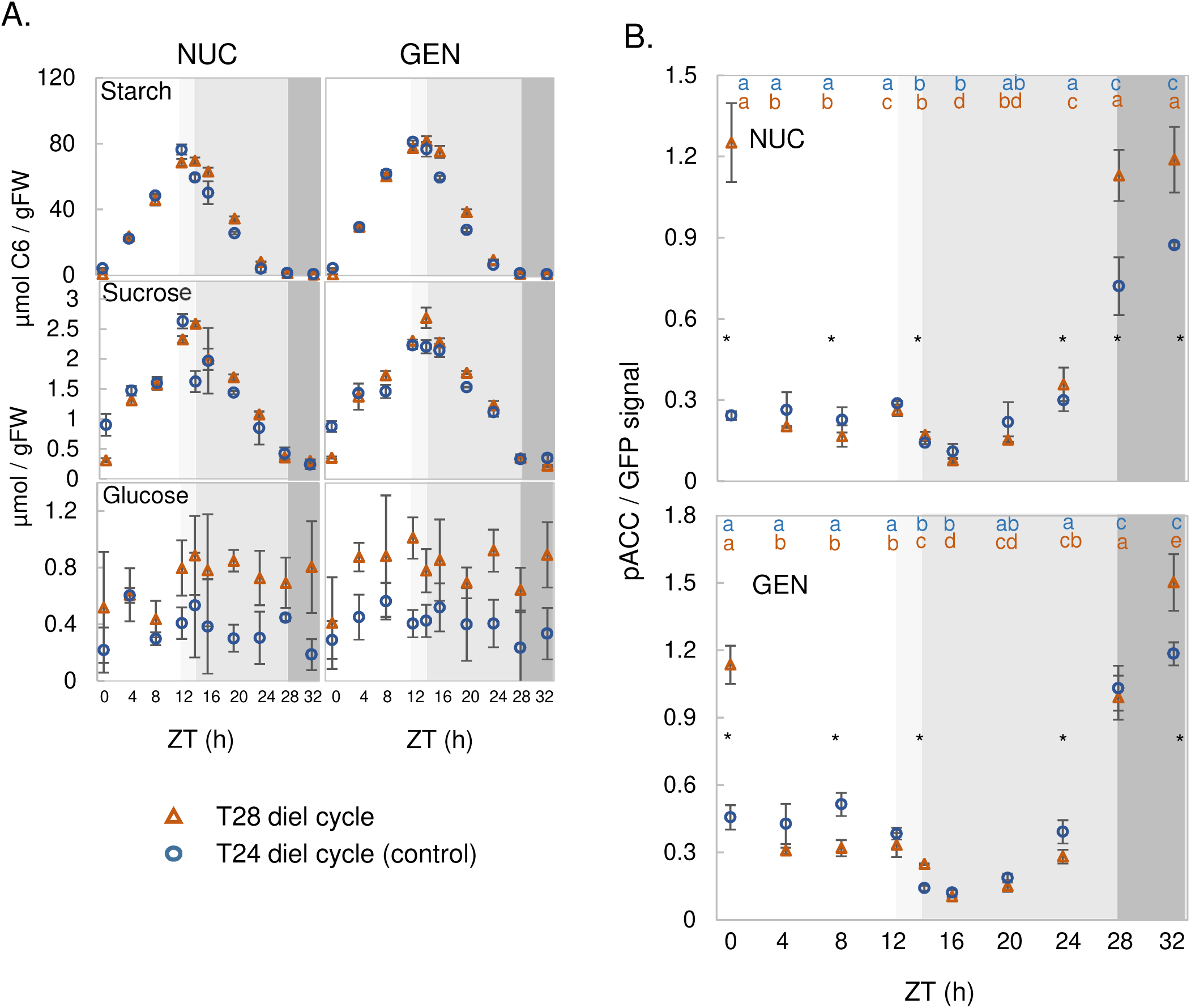
SnRK1 activity in Arabidopsis plants grown in T24 and T28 light-dark cycles. NUC and GEN lines were grown from sowing in a 14-h light / 14 h dark (T28) cycle under 160 µmol m-^2^ s^-1^ irradiance for 19 days, in parallel with control plants that were grown in a 12-h light / 12-h dark (T24) cycle. At 20 DAS (as defined by the T24 cycle), starting at dawn, plants were harvested at 4-h intervals throughout the light-dark cycle (open symbols). At the end of the light-dark cycle, additional plants were left in darkness and harvested for two further time points for the plants grown in a T24 cycle (i.e. at ZT28 and ZT32), and for one further time point for plants grown in a T28 cycle (i.e. at ZT32). (A) Soluble sugars and starch. (B) Phosphorylation of NUC (top) and GEN (bottom) SnRK1 reporter polypeptides. The white background represents light in both T cycles and the grey shaded areas represent darkness (ZT12-ZT24 for the T24 cycle and ZT14-ZT28 for the T28 cycle) in one or both T cycles: pale grey denotes darkness in the T24 and light in the T28 cycle, mid-grey denotes darkness in both T-cycles, and dark grey represents extended night in both T cycles. Black closed symbols represent time points in extended darkness. Results are shown as mean ± S.D. (n = 3 biological replicates). Statistical analysis: letters indicate significant (*P*<0.05) changes between different times for a given genotype and asterisks represent significant (*P*<0.05) differences between T28 vs T24 at a given time point according to a one-way ANOVA with pairwise multiple comparison post-testing using the Holm-Sidak method. The original and additional data are available in Supplementary Dataset S1-Exp.5. The control time-series is also shown in Figure 1 (grey symbols).

### Manipulation of Tre6P levels

Tre6P is a signal of sucrose levels (Lunn *et al*., 2006; 2014; Figueroa and Lunn, 2016; Fichtner and Lunn, 2021) and is widely accepted as an inhibitor of SnRK1, at least *in vitro* (Zhang *et al*., 2009; Zhai *et al*., 2018). However, the evidence that these interactions occur *in vivo* is largely correlative (Martínez-Barajas *et al*., 2011; Wingler *et al*., 2012; Nunes *et al*., 2013; Henry *et al*., 2015; Nuccio *et al*., 2015; Griffiths *et al*., 2016; Peixoto *et al*., 2021; Zacharaki *et al*., 2022). We exploited our SnRK1 reporter lines to develop a genetic approach to test whether Tre6P inhibits SnRK1 activity *in vivo*. We have previously used inducible-TPS (iTPS) lines, expressing the *Escherichia coli* TPS under the control of the *Aspergillus nidulans* ethanol-inducible ALCOHOL DEHYDROGENASE REGULATOR (alcR) and alcA promoter to investigate the impact of a short-term induced increase in Tre6P on starch, organic acid and nitrogen metabolism (Martins et al.,2013; Figueroa et al., 2016). We now introgressed the NUC and GEN reporters into a well characterized iTPS line (TPS29.2; Martins et al., 2013) and into its corresponding empty-vector control, alcR. Four homozygous lines were generated: two lines containing both the alcR gene and the inducible *TPS* (NUC х iTPS #6.3 and GEN х iTPS #2.4), and two control lines with only the alcR gene (NUC х alcR #1.2 and GEN х alcR #1.1).

These lines were grown in long-day conditions (16-h photoperiod, 160 µmol m^-2^ s^-1^ irradiance). On day 21 after sowing, plants were sprayed at ZT1 with either water (control) or 2% (v/v) ethanol to induce the expression of the bacterial TPS (Figure S6). Two whole rosettes were harvested and pooled at 4, 6 and 8h after spraying. Transient expression of the bacterial TPS protein was detectable within 4 h of induction (Supplementary Figure S6B), consistent with previous experiments with the parental iTPS line (Martins *et al*., 2013; Figueroa *et al*., 2016). After ethanol induction, there was a strong and consistent decrease in SnRK1 nuclear activity (in NUC x iTPS) and a small decrease in overall SnRK1 activity after 4 and 6 but not after 10-h in GEN x iTPS (Supplementary Figure S6A; Supplementary Dataset S1 – Exp.6). This led us to focus on the NUC х iTPS line (#6.3) in subsequent experiments. There was no consistent effect of ethanol on NUC or GEN phosphorylation in the control alcR lines, in which alcR is expressed but there is no bacterial *TPS* gene to be expressed under the control of the alcA promoter. This control excludes any risk of off-target effects in the presence of ethanol or due to expression of alcR (Randall, 2021).

To confirm the inhibitory effect of Tre6P induction in the light period, we conducted a more extensive experiment. Line #6.3 was grown in equinoctial conditions under 160 µmol m^-2^ s^-1^ irradiance for 20 days (the standard conditions of previous experiments), sprayed on 21 DAS with either water or 2% (v/v) ethanol at ZT4, and triplicate samples were harvested at ZT7, 9 and 11 and separately analysed (Figure 5; Supplementary Dataset S1 – Exp.7). Ethanol spraying led to a 41, 169 and 130% increase in Tre6P levels at ZT7, ZT9 and ZT11, compared to control plants (Figure 5). Starch accumulation was marginally enhanced, sucrose levels decreased at ZT7 and ZT11 (Figure 5), reducing sugars were not consistently or significantly affected, and the levels of many organic acids increased (Supplementary Figure S7; Supplementary Dataset S1 – Exp.7), as previously reported by Figueroa *et al*. (2016). NUC phosphorylation showed a significant but small decrease (12-23%) following ethanol spraying, compared to the water sprayed control (Figure 5). This provides evidence that Tre6P indeed inhibits nuclear SnRK1 activity *in vivo*, but the impact seems to be rather small at the whole-rosette level.

**Figure 5:**
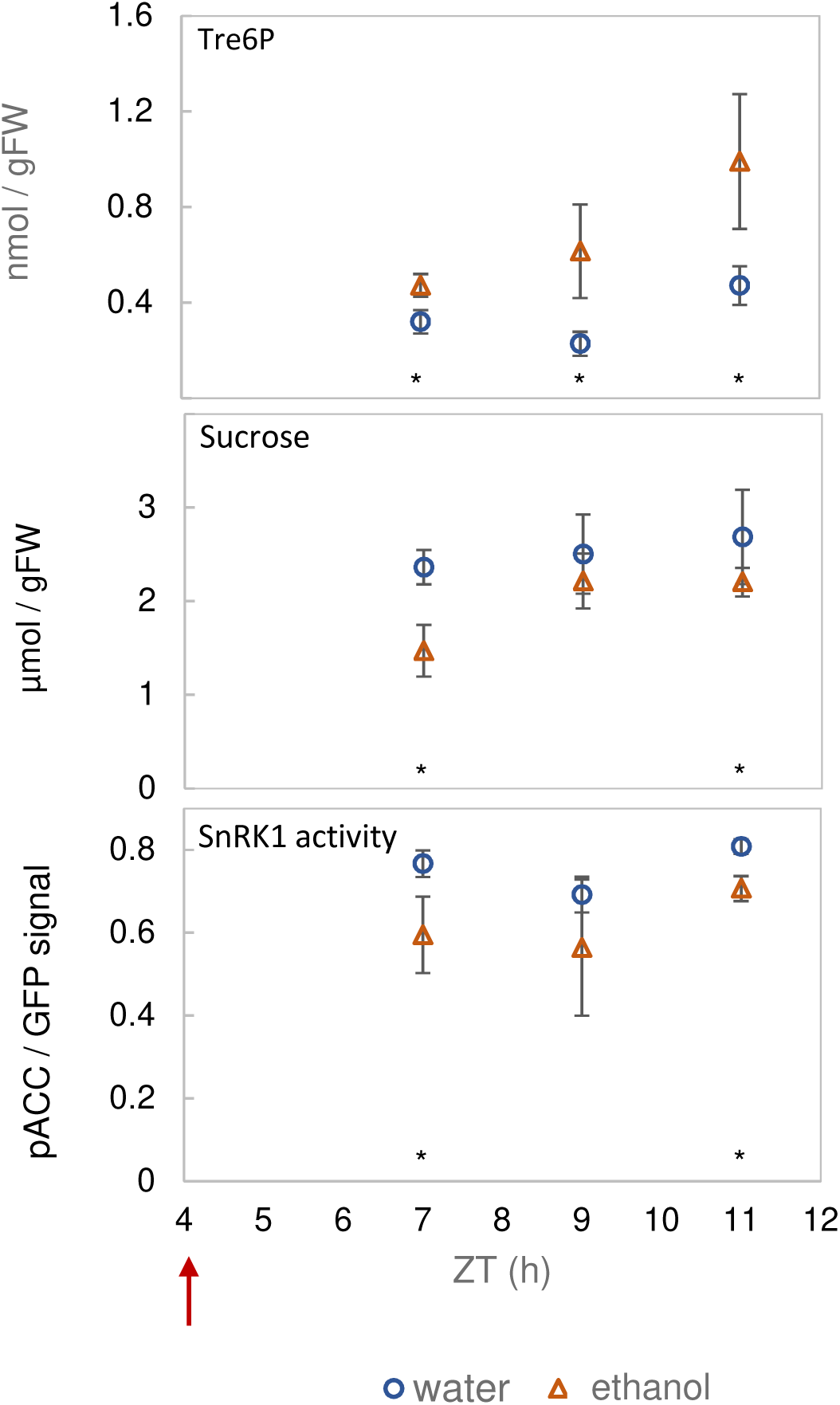
Elevated Tre6P in the light period inhibits nuclear SnRK1 activity. Line #6.3 (NUC х iTPS) was grown in a 12-h photoperiod at 160 µmol m^-2^ s^-1^ irradiance for 20 days. At 21 DAS, plants were sprayed at ZT4 (red arrow) with 2% (v/v) ethanol to induce the expression of bacterial TPS (otsA) or with water as a mock-induction control, and then harvested at ZT7, ZT9 and ZT11 for measurement of Tre6P (top panel), sucrose (middle) and the phosphorylation status of the NUC polypeptide (bottom) as a readout of nuclear SnRK1 activity *in vivo*. Results are shown as mean ± S.D. (n = 3-4 biological replicates). Statistical analysis: asterisks indicate significant (*P*<0.05) differences between the ethanol-induced plants and water controls according to a one-way ANOVA with pairwise multiple comparison post-testing using the Holm-Sidak method. Additional metabolites are shown in Supplementary Figure S7. The original data are available in Supplementary Dataset S1 – Exp.7.

We hypothesized that if Tre6P were not the only regulator of SnRK1, other factors might counteract, and thereby mask, the effect of Tre6P. Therefore, we carried out a further experiment in which TPS was induced in conditions when sugars and other metabolites were low, and might not mask the expected inhibition by Tre6P. Plants were grown in short-day conditions (6-h light / 18-h dark) for 21 DAS with 160 µmol m^-2^ s^-1^ irradiance, sprayed with either water or 2% (v/v) ethanol towards the end of the night (ZT20) and harvested at ZT20, 22, 24, 25 and 27 (Figure 6). We included an alcR control line, and all comparisons were made between ethanol-induced NUC х iTPS (#6.3) and ethanol-induced NUC х alcR #1.2 as the control. Under these conditions, starch was almost completely exhausted and soluble sugars were at low levels (Supplementary Figure S8A). Tre6P levels declined strongly between ZT20 and ZT27 in the control. Induction of TPS slowed down this decline (Figure 6), with Tre6P levels at ZT24, 25 and 27 being significantly higher in ethanol-sprayed NUC х iTPS #6.3 than in the NUC х alcR (#1.2) control at the same time (by 59, 112 and 33%, respectively). Phosphorylation of NUC in the control rose between ZT20 and ZT24, and rose further at ZT25 and ZT27 (Figure 6). This resembles the response of wild-type plants at the end of the night and in an extended night (see above). Unexpectedly, induction of TPS did not attenuate but instead promoted this increase in nuclear SnRK1 activity.

**Figure 6:**
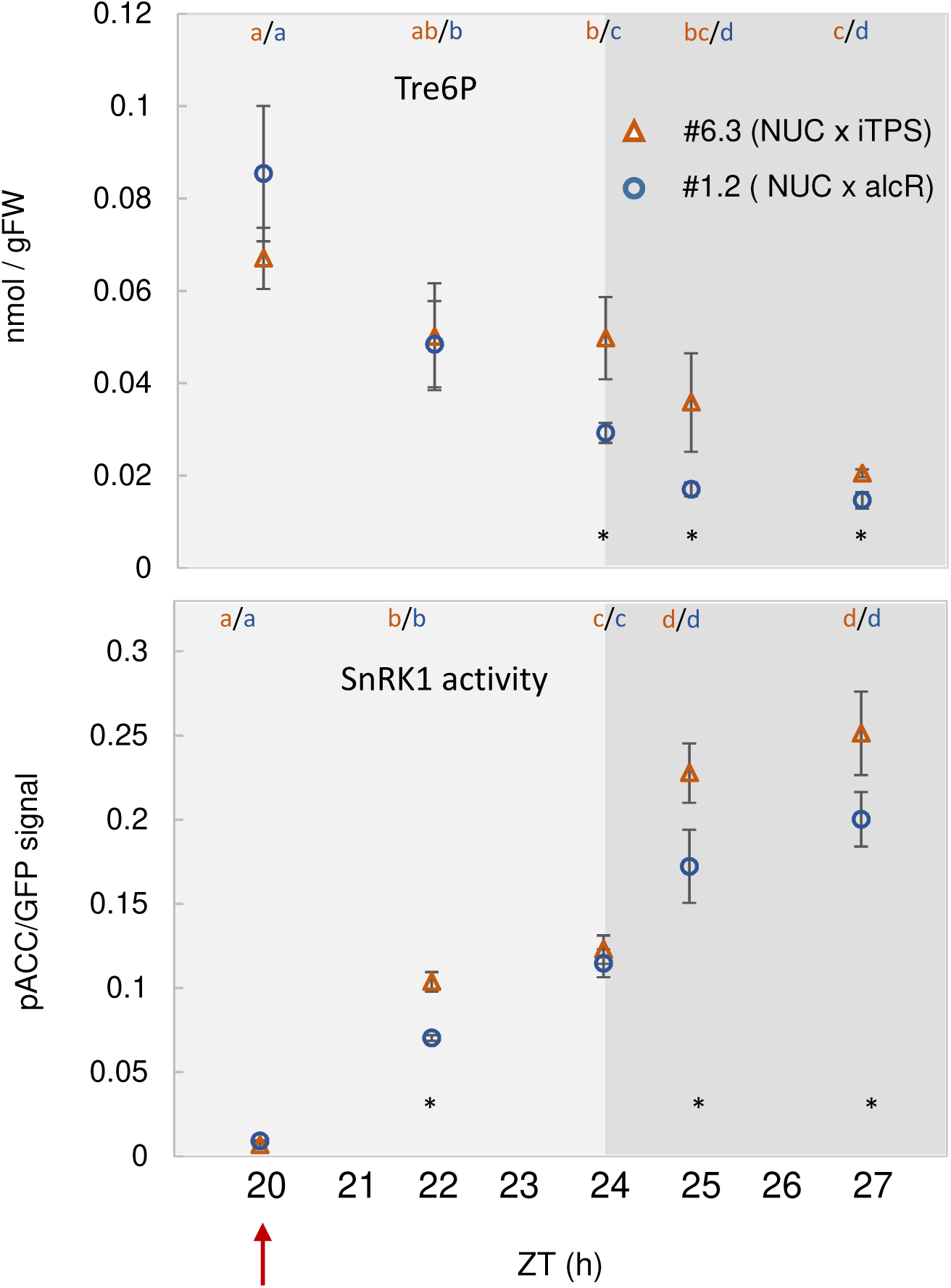
Transiently high Tre6P at the end of the night leads to an increase in nuclear SnRK1 activity. Line #6.3 (NUC х iTPS) and #1.2 control (NUC х alcR) were grown in short-day conditions (6 h light / 18 h dark) at 160 µmol m^-2^ s^-1^ irradiance for 25 days. At 26 DAS, plants were sprayed with 2% (v/v) ethanol at ZT20 (red arrow) to induce the expression of bacterial TPS (otsA). Rosettes were harvested at ZT20, ZT22 and ZT24 (end of normal night), and at ZT25 and ZT27 (extended darkness)for measurement of: Tre6P (top panel) and the phosphorylation of the NUC polypeptide (bottom) as a readout of nuclear SnRK1 activity *in vivo*. Results are shown as mean ± S.D. (n=3-4 biological replicates). Statistical analysis: letters indicate significant (*P*<0.05) changes between different times for a given genotype and asterisks indicate significant (*P*<0.05) differences between the #6.3 (NUC х iTPS) and #1.2 control (NUC х alcR) lines according to a one-way ANOVA with pairwise multiple comparison post-testing using the Holm-Sidak method. Additional metabolites are shown in Supplementary Figure S8. The original data are available in Supplementary Dataset S1 – Exp.7.

The increase in Tre6P levels was accompanied by a slight slowing of starch breakdown and lower glucose levels (Supplementary Figure S8A). This confirms the response previously reported in Martins *et al*. (2013), but in the present case for induction at a later time in the night. The increase in Tre6P levels was also accompanied by increases in malate, pyruvate and glycerol 3-phosphate (Gly3P) and a decrease in phospho*enol*pyruvate (PEP) levels (Supplementary Figure S8B; Supplementary Dataset S1 – Exp.8), indicating that the stimulation by Tre6P of anaplerotic flux that was previously reported in the light (Figueroa *et al*., 2016) may also occur at the end of the night. Some of these downstream responses to elevated Tre6P may lead to the unexpected increase in nuclear SnRK1 activity after iTPS induction.

Overall, these results show that Tre6P is not the sole regulator of SnRK1, and its inhibitory effect appears to be context dependent. Indeed, in some conditions, secondary changes caused by Tre6P induction may lead, indirectly, to a stronger positive effect on SnRK1 activity than the direct inhibitory effect of elevated Tre6P itself.

### Correlation analysis of relationships between SnRK1 activity and metabolite levels

Taken together, our experiments point to complex and somewhat flexible regulation of SnRK1 activity in conditions where C status is changing in a range above that prevailing during starvation. We therefore investigated more systematically the relationship between diel changes of metabolite levels and NUC or GEN phosphorylation.

We did this first in the five control experiments in equinoctial conditions of Figure 1. We performed regression analysis separately for each experiment on all biological replicates and time points across the diel cycle (Table I, for plots see Supplementary Dataset S2). Tre6P levels were not correlated with either NUC (R^2^ = 0.007, 0.004 and 0.004) or GEN phosphorylation (R^2^ = 0.001 and 0.046). Glc6P levels were not consistently related to NUC phosphorylation with R^2^ values of 0.048, 0.21 (negative and significant, P = 0.022) and 0.02 but were negatively correlated with GEN phosphorylation with R^2^ values of 0.22 (P = 0.03) and 0.58 (P = 2.24×10^-5^). Similar relationships were observed for Glc1P which, due to the near-equilibrium reaction catalysed by phosphoglucomutase, can be expected to change in parallel with Glc6P. Starch was not correlated with either NUC or GEN phosphorylation. Sucrose was not correlated with NUC and was negatively correlated with GEN phosphorylation in one but not the other experiment. Glucose was not correlated with NUC, and positively correlated with GEN phosphorylation in one but not the other experiment.

**Table 1:** Correlation between metabolite levels and NUC or GEN phosphorylation. Presented are R^2^ and p-values of linear regression, performed separately for the three independent control experiments compiled in Figure 1. Linear regression analysis was performed with the “least squares” method, using individual replicates for the complete diel cycle (ZT0 to ZT24). The column +/- provides information about the slope of the regression: positive (+), negative (-). Significant correlations (p < 0.05) are highlighted in blue (positive) or red (negative). n.d. not determined. Individual plots are provided in Supplementary Dataset S2.

| Experiment | Metabolite | NUC |  |  | GEN |  |  |
| --- | --- | --- | --- | --- | --- | --- | --- |
|  |  | R<sup>2</sup> | p-value | +/- | R<sup>2</sup> | p-value | +/- |
| 1 | Tre6P | 0.007 | 0.673 | + | 0.001 | 0.965 | + |
|  | Glc6P | 0.048 | 0.270 | - | 0.225 | 0.030 | - |
|  | Glc1P | 0.056 | 0.236 | - | 0.036 | 0.412 | - |
|  | Starch | 0.006 | 0.706 | + | 0.051 | 0.324 | - |
|  | Sucrose | 0.085 | 0.139 | - | 0.058 | 0.295 | - |
|  | Glucose | 0.121 | 0.076 | + | 0.272 | 0.015 | + |
| 2 | Tre6P | 0.004 | 0.778 | + | 0.046 | 0.325 | - |
|  | Glc6P | 0.215 | 0.022 | - | 0.583 | 2.24x10<sup>-5</sup> | - |
|  | Glc1P | 0.418 | 0.001 | - | 0.683 | 1.17x10<sup>-6</sup> | - |
|  | Starch | 0.081 | 0.190 | - | 0.081 | 0.189 | - |
|  | Sucrose | 1x10<sup>-8</sup> | 0.685 | - | 0.324 | 0.005 | - |
|  | Glucose | 0.054 | 0.687 | - | 2x10<sup>-5</sup> | 0.985 | - |
| 3 | Tre6P | 0.004 | 0.233 | - | n.d. | n.d. |  |
|  | Glc6P | 0.002 | 0.766 | - | n.d. | n.d. |  |
|  | Glc1P | 0.02 | 0.373 | + | n.d. | n.d. |  |
|  | Starch | 0.032 | 0.364 | - | n.d. | n.d. |  |
|  | Sucrose | 0.007 | 0.678 | + | n.d. | n.d. |  |
|  | Glucose | 0.013 | 0.561 | - | n.d. | n.d. |  |

We also performed regression analysis separately on the light period (ZT4-12) and night (ZT16-24) datasets (Supplementary Table S1). There were almost no significant relationships in the light period. This can be expected as even though the levels of most sugars, sugar-phosphates and many other metabolites change, SnRK1 activity is rather constant during the light period. During the night there were many negative relationships between NUC/GEN phosphorylation and many metabolites, including Tre6P, Glc6P, Glc1P, starch and sucrose. These negative relationships often reflect falling levels of many metabolites towards the end of the night, as starch is gradually exhausted. They may individually or collectively contribute to the rise in NUC and GEN activity at this time.

Pairwise correlation analyses between metabolites (Supplementary Table S2) revealed that most of the measured metabolites are positively correlated with each other. This reflects the highly coordinated nature of the changes in metabolism in diel cycles, and underlines that some of the observed correlations between metabolites and NUC or GEN phosphorylation may be secondary.

The diel changes in SnRK1 activity and metabolite levels include abrupt changes after illumination or darkening. These may not be well represented in correlations based on changes across the entire cycle. To place more weight on such transient changes, we calculated the difference between the average absolute levels at consecutive time points and divided this by the time difference to obtain the direction and rate of change of a given trait in a given time interval. Compared to absolute values, derivative values provide more information about changes in the balance in metabolism. These derivative values were used for linear regression analyses as described above, except that analyses were only performed for the entire diel cycle (Supplementary Table S3; Supplementary Dataset S2). Changes in both NUC and GEN phosphorylation are significantly and negatively correlated to changes in Glc6P and Glc1P, with changes in Glc6P showing the strongest reciprocal agreement with changes in GEN (highest R^2^ and lowest p-values).

We also performed correlation analyses in the experiments where plants were transferred to continuous light, or grown in T28 cycles, or in which iTPS lines were used to elevate Tre6P levels (Supplementary Dataset S2; Supplementary Tables S4-5, Supplementary Text S1). These analyses again detected correlations between NUC or GEN phosphorylation and Glc6P and Glc1P, and also with Tre6P. Further, in the experiments in which Tre6P was induced in the light period, (Fig 5), NUC phosphorylation correlated negatively with Tre6P in the control plants (R^2^ = 0.67, p = 0.76) but not when ethanol was sprayed to increase Tre6P (R^2^ = 0.12, p = 0.004) (Supplementary Table S6). This is consistent with the idea that SnRK1 activity is regulated by a network including Tre6P and that this network adjusts to decrease the sensitivity to Tre6P when Tre6P levels are artificially elevated.

## Discussion

### Monitoring in vivo SnRK1 activity in plants by expression of a chimeric protein reporter

Two approaches have been widely utilized to study the regulation of SnRK1 activity in plants. One is *in-vitro* phosphorylation assays using a peptide with a conserved SNF1/SnRK1/AMPK recognition sequence (Dale *et al*., 1995). *In-vitro* studies are useful for identifying potential ligands and regulators, including interacting proteins, and can reveal changes in maximum catalytic activity of SnRK1. However, removal of the enzyme from its *in-vivo* context means that activity measurements on *in-vitro* extracts provide only a limited information about activities *in vivo* The other approach is to monitor *in vivo* readouts of SnRK1 activity, such as changes in the abundance of marker gene transcripts (Baena-González *et al*., 2007; Zhang *et al*., 2009; Ramon *et al*., 2019; Crepin and Rolland, 2019). However, while relatively easy to quantify, transcripts are only indirect readouts of SnRK1 activity. Their abundance reflects changes in the activity of transcriptional regulators that are the direct targets for phosphorylation by SnRK1, but can also be influenced by SnRK1-independent changes in transcriptional or post-transcriptional regulation. The ACC1-based polypeptide reporters developed by Deroover *et al*. (2016) have been used to monitor *in-vivo* SnRK1 activities in Arabidopsis (Sanagi et al., 2021; Muralidhara et al., 2021; Henninger et al., 2021; Belda-Palazon et al., 2022), and we applied this approach to study *in-vivo* SnRK1 activity during the diel cycle in Arabidopsis rosettes. This approach does have some potential limitations. The ACC1-based polypeptide might be phosphorylated by other protein kinases in addition to SnRK1, or the extent of phosphorylation may be affected by changes in the activity of endogenous protein phosphatases. Further, the approach detects phosphorylation of a heterologous polypeptide, and so might not capture modifications of SnRK1 function that alter its ability to recognize different endogenous protein targets. In addition to the tests performed in leaf mesophyll protoplasts by Sanagi et al. (2021), we validated our approach by confirming that phosphorylation of the reporter increased when SnRK1 activity *in planta* was enhanced by extended darkness (Supplementary Figure S3A) or overexpression of SnRK1α1, and that phosphorylation of the reporter decreased when SnRK1 activity was suppressed by expression of a dominant negative form (35S:SnRK1α1^K48M^; Supplementary Figure S3B). Nevertheless, the potential complications noted above should be borne in mind when interpreting the results.

SnRK1 protein is located in the nucleus and in various extra-nuclear locations, and probably has different targets in each of these cellular locations (see Introduction and Crozet *et al*., 2014; Ruiz-Gayosso *et al*., 2018; Ramon *et al*., 2019; Baena-González and Lunn, 2020). We introduced GFP-tagged reporter constructs with a nuclear localization sequence (NUC) and without a nuclear localization sequence (GEN), and confirmed that the former is localized to the nucleus, whereas the latter has a more general location inside and outside the nucleus (Supplementary Figure S2). In many treatments, including unperturbed diel cycles (Figure 1) and after transfer to extended darkness (Supplementary Figure S3A), the phosphorylation state of the NUC and GEN peptides changed in a qualitatively similar manner. However, in some perturbations they changed in a dissimilar or even opposite manner (see Figures 2-3), presumably reflecting differing or even opposing changes in SnRK1 activity inside and outside the nucleus, which might be due to differential regulation or to altered subcellular distribution of SnRK1.

### Diel changes in SnRK1 activity point to it operating in a network that buffers C status

It is well known that SnRK1 activity increases in stress and starvation conditions (Baena-Gonzalez *et al*., 2007; Baena-Gonzalez and Sheen, 2008; Mair *et al*., 2015; Cho et al., 2016; Pedrotti *et al*., 2018; Ramon *et al*., 2019). However, it has emerged that SnRK1 also plays a role in benign conditions (Ramon *et al*., 2019; Baena-González and Lunn, 2020; Peixoto *et al*., 2021). After establishing that our *in vivo* assay captures the expected increase in SnRK1 activity in C-depleted conditions, we focused on monitoring SnRK1 activity in conditions where C availability is changing but without stressful extremes of C starvation or surplus. To do this, we grew Arabidopsis in a 12-h photoperiod with 160 µmol m^-2^ s^-1^ irradiance as a standard condition in which growth is restricted by the C supply, but not strongly (Sulpice *et al*., 2014).

One major finding was that *in-vivo* SnRK1 activity was relatively high in the light period, fell after ZT12 (coinciding with darkening in our standard growth conditions) to a minimum at ZT16 and then rose until the end of the night (Figure 1). Activity during the unperturbed diel cycle was several times lower than in extended darkness or other treatments that led to C depletion (see Figure 4, Figure 6, Supplementary Fig 3A). Nevertheless, it was unexpected to observe higher SnRK1 activity in the light period than during the first part of the night, because photosynthesis provides far more C in the light than can be provided by mobilizing C reserves like starch in the night. Plants face a daily challenge to balance the utilization of newly fixed C for immediate growth with the need to lay down C reserves (e.g. starch) and to use these prudently to survive through the night. Therefore, plants must carefully regulate the rate of starch accumulation in the light period, the rate of starch mobilization at night, and the rate at which C is used for growth at different times in the diel cycle (Ishihara *et al*., 2015; Ivakov *et al*., 2017; Mengin *et al*., 2017). The diel pattern of SnRK1 activity that we observed presumably reflects coordinated regulation of C utilization for reserve formation and growth throughout the diel cycle. Incidentally, the relatively low SnRK1 activity during the night adds to the evidence that C homeostasis is maintained across the diel light-dark cycle. In agreement, sucrose levels remained relatively high for much of the night.

A second unexpected finding was that SnRK1 activity did not increase in response to a decrease in overall C availability in the light. Illumination from dawn onwards at a lower irradiance than growth irradiance did not lead to an increase in the phosphorylation status of either the NUC or GEN reporter polypeptides, compared to that at growth irradiance (Figures 2 and 3). Although photosynthesis was not measured in these experiments, the low irradiance treatments did lead to a consistent decrease in the levels of sucrose and reducing sugars (Figures 2B and 3B; Supplementary Dataset S1 – Exp.3-4; see also Moraes e*t al*., 2019). Further, these changes in irradiance occurred in the range where Arabidopsis photosynthesis is light-limited (see Borghi *et al*., 2019), and previous studies have shown that similar drops in irradiance lead to a large decrease in photosynthetic rate (see Fernandez *et al*., 2017; Moraes *et al*., 2019). On the other hand, treatments that led to a decreased dusk starch content and slower rate of starch mobilization during the night did lead to a consistent increase *in-vivo* SnRK1 activity in the first part of the night.

These observations point to SnRK1 being regulated in a more flexible manner in the light than in the dark. This flexibility is also revealed by the response when plants are transferred to low irradiance for one light period and then returned to growth irradiance. This perturbation led to higher levels of sugars and faster accumulation of starch when plants were returned to growth irradiance, compared to control plants that had been left at growth irradiance (Figure 2B). These observations indicate that low irradiance results in a restriction of growth on the following day after returning plants to previous growth irradiance (see also Moraes et al., 2019). SnRK1 activity showed a complex pattern after returning to the previous growth irradiance (Figure 2); phosphorylation of NUC was lower than in control plants, reflecting the higher levels of sugars, whereas phosphorylation of GEN resembled that in control plants implying that extra-nuclear SnRK1 activity was higher than in control plants despite sugar levels also being higher. It is known that growth in the light period is restricted when Arabidopsis experiences a shortfall in C in the preceding 24-h cycle (Gibon et al., 2004; Sulpice *et al*., 2014; Flis et al., 2016; Mengin *et al*., 2017; Moraes *et al*., 2019). Our observations indicate that increased extranuclear SnRK1 activity might contribute to this restriction of growth and the resulting increase in starch accumulation.

There may also be more ways for the plant to cope with a lower rate of photosynthesis in the light than there are to adjust to a shortfall of starch at night. For example, the impact of decreased irradiance on Glc6P levels in the light period was less marked than the impact on sugar levels (Supplementary Figure S4, see also Moraes et al., 2019). This may reflect compensatory responses that buffer the level of this key metabolic intermediate in the light, for example, by decreasing starch synthesis, and by decreasing consumption of hexose phosphates for sucrose synthesis in the cytosol by allosteric and post-transitional inhibition of sucrose-phosphate synthase (Huber and Huber, 1996; Smith and Stitt, 2007; Lunn, 2016).

A third finding was that an endogenous circadian rhythm may contribute to the diel changes in SnRK1 activity, at least in the middle of the 24-h cycle, when SnRK1 activity decreased between about ZT12 and TT16. The decrease after ZT12 was partly retained in low continuous light (Figure 3) and was also observed in plants growing in a T28 cycle, even though they were in the light between ZT12 and ZT14 (Figure 4). The parallel changes in sugars and predicted extra-nuclear SnRK1 activity after returning plants from low to growth irradiance may also point to an endogenous signal that relays information about C status in the previous 24-h cycle.

### SnRK1 activity may be regulated by multiple metabolic inputs including Tre6P and hexose phosphates

The unexpected diel response pointed to SnRK1 operating within a network that controls C utilization in the daytime and at night to maintain C homeostasis. SnRK1 activity integrates various inputs including multiple signals from metabolism (Polge and Thomas, 2007; Baena-González and Sheen, 2008; Wurzinger *et al*., 2018). We asked to what extent diel changes in SnRK1 activity are accompanied by reciprocal changes in the levels of sugars that might be viewed as indicators of C availability, or reciprocal changes in the levels of metabolites, like Tre6P, Glc6P and Glc1P, that are known to inhibit SnRK1 *in vitro* (Toroser *et al*., 2000; Zhang *et al*., 2009; Debast *et al*., 2011; Delatte *et al*., 2011; Piattoni *et al*., 2011; Nunes *et al*., 2013a; Coello and Martínez-Barajas, 2014; Emanuelle *et al*., 2015; Griffiths *et al*., 2016; Bledsoe *et al*., 2017; Henry *et al*., 2020; Baena-González and Lunn, 2020).

The role of Tre6P in regulation of SnRK1 has attracted wide attention (Zhang *et al*., 2009; Nunes *et al*., 2013; Henry *et al*., 2015; Zhang *et al*., 2015; Bledsoe *et al*., 2017; Baena-González and Lunn, 2020). To directly test whether an induced increase in Tre6P leads to inhibition of *in-vivo* SnRK1 activity, we crossed our reporter constructs into lines carrying an inducible bacterial TPS construct. An induced 1.4- to 2-fold increase in Tre6P in the light period indeed led to 12-23% decrease in phosphorylation of the NUC reporter polypeptide (Figure 5), and a weak decrease for the GEN reporter polypeptide (Supplementary Figure S6A). On the other hand, during diel cycles in wild-type plants Tre6P levels did not show a consistent negative relationship to NUC or GEN phosphorylation (Figure 1; Table 1). Whereas Tre6P levels rose 3- to 5-fold during the light period, peaked at ZT12 (ED) and then fell, (Figure 1B), SnRK1 activity did not change during the light period and, after darkening, fell by 2- to 3-fold to a minimum at around ZT16 (Figure 1A). Also, comparison of the magnitude of the changes in Figure 5 and Figure 1 indicates that the diel changes of Tre6P are too small to explain the diel changes in NUC phosphorylation. In the last part of the night, where there was a negative correlation between Tre6P levels and SnRK1 activity (Supplementary Table S1), an induced increase in Tre6P resulted in an increase rather than a decrease in SnRK1 activity (Figure 6). Whilst these findings provide evidence that Tre6P can contribute to the regulation of SnRK1 *in vivo*, they also show that the contribution of Tre6P is context-dependent and that Tre6P is not the only or even the major factor that regulates SnRK1 activity.

The possible role of hexose phosphates like Glc6P and Glc1P in the *in-vivo* regulation of SnRK1 activity has attracted less attention, despite evidence from *in-vitro* assays that they can inhibit SnRK1 (Toroser et al., 2000; Zhang *et al*., 2009; Piattoni *et al*., 2011; Nunes *et al*., 2013). Several features of our study point to them playing a role *in vivo*. First, visual inspection of diel changes under control conditions reveal that Glc6P and Glc1P often change in a reciprocal manner to SnRK1 activity (Figure 1), peaking in the first part of the night at ZT16 at the same time as the minimum in SnRK1 phosphorylation activity. A negative relationship between SnRK1 and Glc6P was also observed when plants were exposed to a sudden low-light day (Figure 2; see also Moraes *et al*., 2019) or transferred to continuous low light (Figure 3; Supplementary Figure S4). Second, there was a significant negative correlation between Glc6P or Glc1P levels and NUC or GEN phosphorylation in some, though not all, full diel cycles (Table 1) and highly consistent negative correlations in the night (Supplementary Table S1). This conclusion was supported by comparison of the rates of change of Glc6P or Glc1P levels and SnRK1 activity at different times in the diel cycle (Supplementary Table S3). Glc6P and Glc1P are linked by the near-equilibrium reaction catalysed by phosphoglucomutase, with both changing in parallel and Glc6P being about 10-fold higher than Glc1P, therefore, the observed negative relationship between SnRK1 and Glc1P might be secondary to the relationship with Glc6P.

The question also arises what endogenous factor might be responsible for the decline in SnRK1 activity between ZT12 and ZT16, which occurred even when plants were not darkened at ZT12 (see above). One possible explanation is that this decline is driven by diel changes in starch turnover. As shown in Fernandez *et al*. (2017) and Ishihara *et al*. (2022), starch mobilization occurs simultaneously with starch synthesis in the light and speeds up with time in the light, leading from about ZT10 onwards to a marked rise in sucrose and related metabolites, including Tre6P. This might contribute to the decline in SnRK1 activity after ZT12 in continuous light. As suggested in Fernandez *et al*. (2017) and Ishihara *et al*. (2022), this gradual rise in the rate of starch mobilization is itself under circadian control. However, it would be premature to exclude the possibility that the circadian clock exerts a more direct influence on SnRK1 activity, mirroring the recent discovery that SnRK1 is involved in transcriptional regulation of the circadian clock (Jeong *et al*., 2015; Sánchez-villarreal *et al*., 2018; Shin *et al*., 2017; Frank *et al*., 2018). A possible link between starch mobilization and SnRK1 activity is also of interest in view of the report that the dark-expressed SnRK1β3 subunit binds maltose, pointing to a possible role in regulating starch degradation or of starch degradation, via maltose, regulating SnRK1 activity (Ruiz-Gayosso *et al*., 2018).

Taken together, our results point to both Tre6P and hexose phosphates contributing to the regulation of SnRK1 *in vivo*. These metabolites provide complementary information about C status. Tre6P provides information about the amount of sucrose (Lunn *et al*., 2006; Figueroa *et al*., 2016; Figueroa and Lunn, 2016; Fichtner *et al*., 2017; Fichtner and Lunn, 2021), which is a key long-distance transport metabolite and often a C-storage molecule (Lunn, 2016). Hexose phosphate levels reflect the balance between several central metabolic pathways, for example, between photosynthetic C fixation and sucrose and starch synthesis in the light, and between starch mobilization, respiration and biosynthesis in the dark. Further, Tre6P and hexose phosphates may have differing subcellular and intercellular compartmentation. TPS1 protein, and by implication Tre6P, is partly localized to the nucleus (Fichtner *et al*., 2020) whereas hexose phosphates are likely to have a more general subcellular distribution with high levels in the cytosol (Stitt *et al*., 1980; Gerhardt *et al*., 1987). Such differences in metabolite compartmentation might underlie some of the differing responses of the NUC and GEN reporters. Further, TPS1, and by implication Tre6P, is very abundant in companion cells and surrounding phloem parenchyma (Fichtner *et al*., 2020) whereas hexose phosphates are probably distributed more generally across cell types, including high levels in photosynthetic mesophyll cells. This may introduce important nuances into regulation of SnRK1 activity, whose detection would require cell-specific expression of the reporter constructs.

### Changes in complex composition might impact on diel SnRK1 activity

An additional regulatory mechanism that might contribute to observed diel changes in SnRK1 activity would be changes in subunit composition, including expression or stability of its subunits, such as by SUMOylation and ubiquitination of SnRK1α1, SnRK1β1, SnRK1β2 SnRK1β3, and by myristoylation of SnRK1β1 and SnRK1β2 (Ruiz-Gayosso *et al*., 2018; Ramon *et al*., 2019; Blanco *et al*., 2019; Wang et al., 2020). These were shown to affect cellular localization, primarily by translocating SnRK1 in and out of the nucleus. We cannot yet be certain whether changes in SnRK1 composition contribute to the diel changes in SnRK1 activity detected using the ACC reporter constructs. In the future it would be of interest to compare changes in SnRK1 composition and SnRK1 location with the relative rates of NUC and GEN phosphorylation.

### Comparison of in-vivo SnRK1 activity with responses of SnRK1 downstream targets highlights potential flexibility in downstream signalling pathways

As already mentioned, downstream target genes identified in protoplast overexpression studies have been widely used to make inferences about SnRK1 activity *in vivo*. We inspected published data to learn if the diel changes in transcript abundance of these genes is consistent with the diel changes of *in-vivo* SnRK1 activity, as monitored by reporter peptide phosphorylation. To do this, we focused on studies in which Arabidopsis was grown in the same conditions as those used in the current study. As shown in Supplementary Figure S9A, which replots data from Usadel *et al*. (2008), there was a fairly consistent pattern with the abundance of SnRK1-induced transcripts falling in the light period and rising during the night, and the abundance of SnRK1-repressed transcripts rising in the light and falling during the night (two exceptions were *AXP* and *EXP10*). This pattern contrasts with the response of the NUC/GEN reporter polypeptide phosphorylation, which was relatively high in the light period and fell in the first part of the night (Figure 1A). In a second comparison, we inspected the response of a set of SnRK1 target transcripts after a sudden decrease in irradiance (Moraes *et al*., 2019), similar to the treatment in Figure 2 and the first 12 h after transfer to continuous low light in Figure 3 (Supplementary Figure S9B). Transfer from 160 to 60 µmol m^-2^ s^-1^ led, for SnRK1-induced genes, to a small but consistent increase in *DIN6* and *BCAT2* and a large increase in *DIN1* transcript abundance and, for SnRK1-repressed genes, to a small but consistent decrease in *CPN60α* and *Asp.P* transcript abundance. This contrasts with NUC/GEN reporter polypeptide phosphorylation, which did not change after transfer to low irradiance (Figures 2-3). In a recent study, Peixoto *et al*. (2021) reported that abundance of SnRK1-induced transcripts was maximal at the end of the night when Tre6P levels were lowest, and minimal at the end of the day when Tre6P levels peaked. However, even though the changes of Tre6P levels in Peixoto *et al*. (2021) resembled those in the current study, the responses of transcript abundance did not match the changes in NUC/GEN reporter polypeptide phosphorylation reported in the current study. Furthermore, whilst transient elevation of Tre6P in an inducible bacterial TPS line led to decreased abundance of transcripts for three SnRK1-induced genes (*BGAL, DIN10, PYL5*) (Peixoto *et al*., 2021), consistent with involvement of Tre6P in diel regulation of SnRK1 signalling, it did not decrease transcript abundance for many other SnRK1-induced genes, such as *DIN1*, *DIN6*, *AXP* and *EXP10*.

There are various possible explanations for discrepancies between SnRK1 activity as monitored by phosphorylation of the NUC/GEN reporter polypeptides and SnRK1 activity as inferred from the abundance of downstream transcripts. It is possible that in some cases gene expression is driven by a fraction of the total SnRK1 pool, and that this fraction may remain undetectable when assaying total SnRK1 activities (Baena-González and Lunn, 2020). The discrepancy may also reflect multiple levels of regulation, for example, not only via changes in SnRK1 activity itself but also via inputs that modify the affinity of SnRK1 for different immediate downstream targets or via inputs that potentiate or attenuate different downstream signalling pathways. Such modifying factors might also explain the varying responses of different SnRK1 downstream transcripts within a given experiment (see above, also Griffiths *et al*., 2016), as well as between different studies. Indeed, of the top SnRK1 target genes, only a few changed in a reproducible manner following SnRK1 induction in protoplasts versus whole plants (Baena-González *et al*., 2007; Ramon *et al*., 2019; Peixoto *et al*., 2021). For example, *DIN1*, *DIN6* and *CAT1*, whose expression is commonly used as a readout of SnRK1 activity, are strongly upregulated in response to transient expression of SnRK1α1 in protoplasts (Baena-González *et al*., 2007), but remain unaffected in response to constitutive overexpression of SnRK1α1 in whole plants (Peixoto *et al*., 2021).

Functionally, operation of SnRK1 in a benign diel cycle as well as during starvation mirrors the differing responses of two sub-sets of C-regulated genes. Many of these show large changes in expression during a regular equinoctial diel cycle (Smith and Stitt, 2007; Usadel *et al*., 2008; Cookson *et al*., 2016, Flis et al., 2016). These genes may contribute to modifications of metabolism and slowing down of growth, and serve to maintain C homeostasis and avoid transient periods of C starvation and the resulting wasteful alternation between growth and catabolism (Ishihara et al., 2015, 2017). Other C-regulated genes do not show changes in expression until starch is exhausted, and may serve to orchestrate catabolic responses in conditions when the plant can no longer avoid C starvation.

In conclusion, by employing an *in-vivo* reporter polypeptide assay we have uncovered unexpected changes in SnRK1 activity under benign light-dark cycles. Continued SnRK1 activity in the light period may contribute to maintenance of diel sugar homeostasis, by restricting C utilization for growth and supporting build-up of C reserves. Further, whilst demonstrating that Tre6P can inhibit SnRK1 activity *in vivo*, we also show that its contribution is context dependent and that hexose phosphates, and probably also other factors, interact to regulate SnRK1 activity and signalling.

## Materials and Methods

### Plant material

All Arabidopsis (*Arabidopsis thaliana* [L.] Heynh.) genotypes were in the Columbia-0 (Col-0) background. Lines expressing the NUC or GEN SnRK1 reporters were generated as described in Sanagi *et al*. (2021). The reporters comprise a 57-amino acid peptide derived from rat (*Rattus norvegicus*) acetyl CoA carboxylase (rACC1), containing the AMPK recognition motif and Ser79 phosphorylation site, fused at the C-terminus to eGFP and a double haemagglutinin tag (Supplementary Figure S1; Deroover *et al*., 2016). The NUC reporter includes an N-terminal SV40 nuclear localization signal that targets the reporter specifically to the nucleus, while the GEN reporter has no added nuclear localization signal and is localized in both the nucleus and cytoplasm (Supplementary Figure S2). The Arabidopsis (Col-0) ethanol-inducible TPS-overexpressor (iTPS 29.2) and empty-vector control AlcR lines were the same as those described in Martins *et al*., 2013. The Arabidopsis (Col-0) 35S:SnRK1α1 over-expression line (Jossier *et al*., 2009) and the 35S:SnRK1α1^K48M^ line, expressing the SnRK1α1 subunit with a mutation that disrupts ATP binding (Cho *et al*., 2016), were kindly provided by Elena Baena-González (Instituto Gulbenkian de Ciência, Oeiras, Portugal) and Hsing-Yi Cho (National Chung-Hsing University, Academia Sinica, Taiwan), respectively.

### Introgression of NUC and GEN SnRK1 reporters

The NUC and GEN SnRK1 reporters were introgressed into established lines with altered SnRK1 activity or inducible expression of TPS by crossing. The respective homozygous parental lines were grown in soil for 4-5 weeks (see below). Pollen from the parental parent was transferred by rubbing anthers from mature flowers against exposed stigmas of emasculated maternal plants under a binocular microscope, and the inflorescence meristems and unopened flower buds were removed. After crossing, the maternal plants were grown on, with regular watering, for 2-3 weeks until siliques had fully developed and started to dry. Siliques were collected, left to dry for a further 3-4 days at room temperature and then kept at 10 °C until seeds were germinated on selective media. After surface sterilization by treatment with 2% (w/v) sodium hypochlorite for 5 min at room temperature and washing in five changes of water, seeds were placed on 0.5x Murashige and Skoog medium (Murashige and Skoog, 1962) containing 1% (w/v) sucrose and appropriate antibiotics for selection of the respective transgenes from each parent: NUC/GEN phosphinothricin (Basta); 35S:SnRK1α1, kanamycin; 35S: SnRK1α1^K48M^, hygromycin; iTPS/alcR; kanamycin. The seeds were stratified for 2 days at 4°C in the dark and then transferred to 22°C in continuous light at 100 µmol m^-2^ s^-1^. After 7 days, surviving seedlings were transferred to soil. The F1 plants were selfed and F2 progeny were screened on selective media, as above, to identify lines that exhibit the expected susceptible: resistant ratio of 7:9. Individual F2 plants were selfed and the F3 progeny were screened on selective media, as above, to identify lines that were homozygous for both transgenic loci.

### Plant growth and harvest

Seeds were sown on a 1:1 mixture of compost (Stender AG, Schermbeck, Germany; https://www.stender.de) and vermiculite in 6-cm diameter pots, covered and kept at 4°C in the dark for 48 h then transferred to a controlled environment chamber (Percival E-36 L chamber, CLF Plant Climatics GmbH, Weringen, Germany; https://www.plantclimatics.de/) with a 16-h photoperiod (160 µmol m^-2^ s^-1^ irradiance provided by white LEDs) and day/night temperatures of 21°C/19°C. After germination, seedlings were transferred to 10-cm diameter pots (four or five seedlings per pot), and grown as above or as described for individual experiments. Whole rosettes were harvested *in situ* and immediately quenched in liquid nitrogen, with four or five plants from the same pot being pooled for each biological replicate. Frozen plant material was ground to a fine powder and stored at -80°C until use.

### Metabolite extraction and analysis

Soluble sugars and starch were extracted and assayed enzymatically as described in Gibon *et al*. (2002) with three-four biological replicates per sample type. For each biological replicate, values were calculated as the average of two measurements. Tre6P, other phosphorylated intermediates and organic acids were extracted with chloroform-methanol and quantified by anion-exchange high-performance liquid chromatography coupled to tandem mass spectrometry as described in Lunn *et al*. (2006), with modifications as described in Figueroa *et al*. (2016).

### Ethanol-inducible iTPS experiments

After introgression of the NUC/GEN SnRK1 reporters, ethanol-inducible TPS (iTPS) and control alcR lines (Martins *et al*., 2013) were grown as described above. On the day of harvest, plants were sprayed with either water (mock induction control) or with 2% (v/v) ethanol to induce expression of the bacterial TPS and thereby increase Tre6P levels. Samples were collected and harvested as described above.

### SDS-PAGE and immunoblot assay of SnRK1 reporter phosphorylation

Proteins were extracted from aliquots (10 mg FW) of frozen, finely powdered plant material by suspension in 100 µl of pre-warmed (90°C) sample buffer (66 mM Tris-HCl, pH 6.8, 10% (v/v) glycerol, 2% (w/v) sodium dodecyl sulphate, 0.13% (w/v) bromophenol blue) quickly vortex mixed, incubated at 90°C for 5 min, then chilled on ice and centrifuged at 13,500 x *g* (4°C) for 3 min. Aliquots of the supernatant containing approx. 12 µg of soluble protein were electrophoresed on a 12% (w/v) polyacrylamide-SDS gel. Proteins were electro-blotted onto nitrocellulose membrane, visualized by staining with 0.2% (w/v) Ponceau Red in 1% (v/v) acetic acid and imaged for use as a protein-loading reference. After blocking in Tris-buffered saline containing 5% (w/v) milk powder, the membrane was incubated overnight at 4°C with rabbit anti-phospho-acetyl CoA carboxylase (Ser79) antibody (#3661; Cell Signaling Technology Europe BV, Leiden, The Netherlands; https://www.cellsignal.de) at a dilution of 1:1500. After washing, NUC/GEN phosphorylation was detected by incubation with IRDye 800CW goat anti-rabbit IgG (H+L) secondary antibody (Li-COR Biosciences GmbH, Bad Homburg, Germany; www.licor.com) at a dilution of 1:9000 and scanning with an Odyssey Imaging Scanner (Li-COR), and quantified using Image Studio Live 5.2 software (Li-COR). Total NUC/GEN polypeptide abundance was detected by probing duplicate blots with a mouse anti-GFP antibody (#11814460001; Sigma Aldrich (Roche), Darmstadt, Germany; www.sigmaaldrich.com) at a dilution of 1:1500, followed by incubation with IRDye 680RD goat anti-mouse IgG (H+L) secondary antibody (Li-COR) at a dilution of 1:9000, and quantified as above.

To minimize the impact of gel-to-gel variation, aliquots from an extended night sample were loaded onto each gel and the signal from this sample was used to normalize the signals from the other samples on the corresponding immunoblot. Two aliquots of the protein extract from each biological replicate were analyzed on separate gels and, after normalization as described above, the average of the two technical replicates was calculated as the reported value for that biological replicate. The signal ratio pACC/GFP was used to normalize for any differences in reporter protein abundance in the tissue and differences in gel loading. See Supplementary Figure S1 for further details.

### Confocal microscopy

To determine the subcellular localization of the NUC/GEN SnRK1 reporter proteins, GFP was visualized by laser confocal microscopy of 6-day-old seedlings. Imaging was performed on a Leica TCS SP5 microscope (Leica Microsystems, Wetzlar, Germany; https://www.leica-microsystems.com) using a 20x objective. To visualize nuclei, NUC samples were stained with 10 μg/ml propidium iodide (PI) solution for 5 to 10 min. A 488 nm laser was used to excite GFP and 561 nm laser to excite PI and their fluorescence emission was detected in the 524-555 nm and 571-640 nm ranges, respectively. Confocal Z sections were acquired every 0.5 μm.

### Statistical analysis

Technical replicates were always averaged to generate a single value for each biological replicate, statistical analysis was restricted to biological replicates and performed using Sigma-Plot 14.5 software (Systat Software GmbH, Düsseldorf, Germany; http://www.systat.de). Significance of changes in metabolite and SnRK1 activity was tested by one-way ANOVA using a pairwise multiple comparison procedure, with post-hoc testing by the Holm-Sidak method (p<0.05), as described in each figure legend.

## Acknowledgements

We thank Sofie Deroover and Tom Broeckx for help with generation and selection of the NUC and GEN SnRK1 reporter lines and Arun Sampathkumar for help with laser confocal microscopy.

## Author contributions

O.A. performed all experiments, statistically analysed data, and contributed to paper writing; T.A.M contributed to experimental design, harvesting, measurement of metabolites and data analysis. V.M. contributed to harvesting and measurement of metabolites, R.F. performed LC-MS/MS metabolite analysis; F.R. conceived and generated the SnRK1 reporter constructs, M.S. assisted data analysis and contributed to writing the paper. J.E.L. conceived the project, assisted data analysis, and contributed to writing the paper.

## Supplementary Data

**Supplementary Figure S1:** *In vivo* SnRK1 activity assay

**Supplementary Figure S2:** Localization of reporter peptides by confocal microscopy

**Supplementary Figure S3:** Validation of the NUC and GEN constructs in whole Arabidopsis rosettes

**Supplementary Figure S4:** Comparison of different low light treatments (supplementary to Figure 2 and 3)

**Supplementary Figure S5:** T28 and T24 cycles; diel Glc6P levels and comparison of SnRK1 activity excluding time points affected by C starvation (supplementary to Figure 4).

**Supplementary Figure S6:** Response of NUC and GEN phosphorylation in crosses with a line containing inducible bacterial TPS.

**Supplementary Figure S7:** Impact of transiently elevated Tre6P in the light period on the levels of selected metabolites (supplementary to Figure 5).

**Supplementary Figure S8:** Transient elevation of Tre6P at the end of the night slows down starch mobilization and leads to low glucose and high organic acid levels (supplementary to Figure 6)

**Supplementary Figure S9:** Transcript abundance for SnRK1 target genes during a diel cycle, after an extension of the night and after a sudden decrease in light intensity for one day.

**Supplementary Table S1:** Correlations between metabolite levels and NUC or GEN phosphorylation, performed separately for the light period and the night (supplementary to Table 1 and Figure 1).

**Supplementary Table S2:** Correlations between metabolite levels (supplementary to Table 1 and Figure 1)

**Supplementary Table S3:** Correlations between changes in metabolite levels and changes in NUC or GEN phosphorylation (supplementary to Table 1 and Figure 1).

**Supplementary Table S4:** Correlations between metabolite levels and NUC or GEN phosphorylation in growth regimes that differed from the equinoctial T24 cycle (supplementary to Figures 3 and 4).

**Supplementary Table S5:** Summary of correlation analysis between changes in SnRK1 NUC and GEN activity to changes in metabolite levels (supplementary to Figures 3 and 4).

**Supplementary Table S6:** Correlation analysis between metabolite levels and NUC phosphorylation following transient elevation of Tre6P in the light period (supplementary to Figure 5, Supplementary Dataset S2 Exp.7).

**Supplementary Text S1:** Correlations between in vivo SnRK1 activity and metabolites in continuous light, in T28 cycles and in experiments where TPS is induced (supplementary to Supplementary Tables S4-S6)

**Supplementary Dataset S1:** *In-vivo* SnRK1 activity and metabolite levels.

**Supplementary Dataset S2:** Regressions between *in-vivo* SnRK1 activity and metabolite levels.

