## Supplementary Figures S1-S9 for "Diel fluctuations in *in-vivo* SnRK1 activity in Arabidopsis rosettes during light-dark cycles"

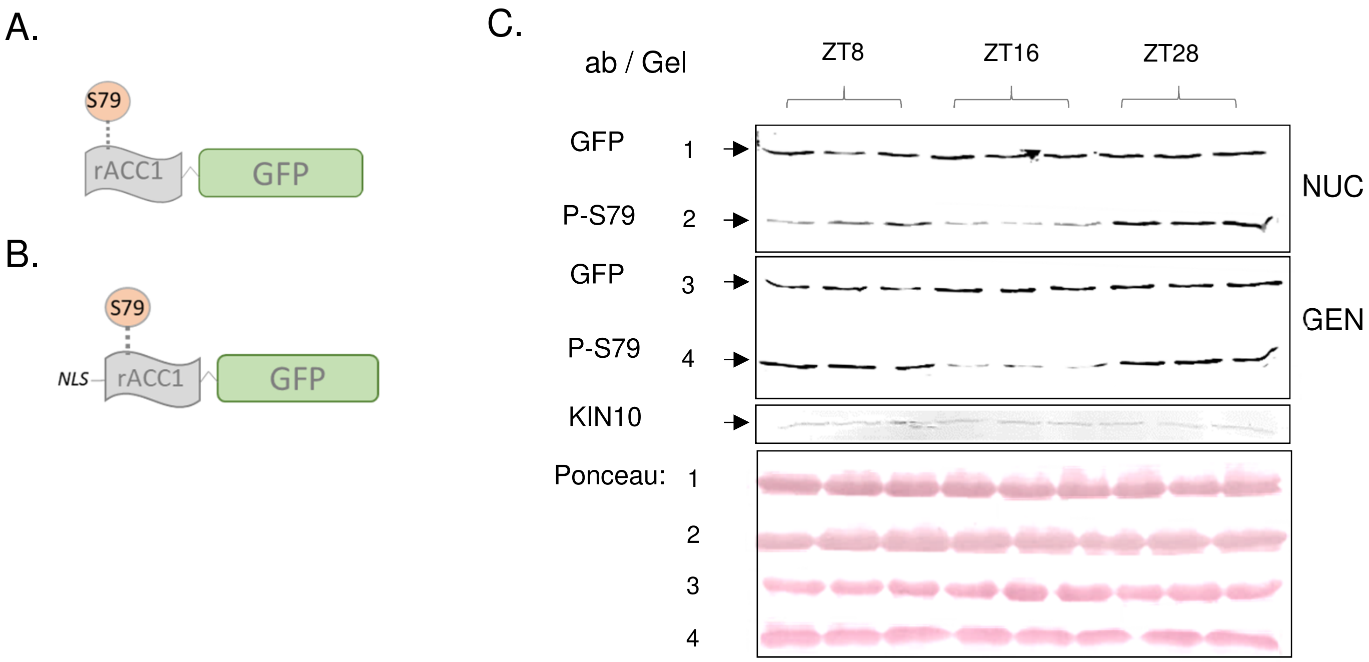


**Supplementary Figure S1: In vivo SnRK1 activity assay.** The homozygous transgenic SnRK1 reporter lines were generated by Sanagi et al. (2021). The principle is based on sequence recognition homology between the three orthologous kinases: mammalian AMPK, plant SnRK1 and yeast SNF1, as described in Deroover *et al.* (2016). In brief, a 57-aa peptide surrounding the Ser79 phosphorylation site of rat acetyl-CoA carboxylase 1 (rACC1) was fused to GFP, cloned into the pCB302 expression vector and transformed into *Arabidopsis thaliana* (Col-0). Two constructs were used: (A) a general reporter (GEN) and (B) a nuclear (NUC) reporter that includes the SV40 nuclear localization signal (see Figure S2 for localization data). (C) Quantification of SnRK1 activity: plants were grown in equinoctial growth conditions (12-h light / 12-h darkness) with 160 µmol $m^{-2}$ $s^{-1}$ irradiance (as described in Methods). Whole rosettes were harvested at the indicated times and immediately quenched in liquid nitrogen. Three biological replicates were harvested at each time point (four or five plants from the same pot were pooled for each biological replicate). About 12 µg of soluble proteins from each replicate were electrophoresed on an SDS-polyacrylamide gel (12% w/v), electro-blotted onto a nitrocellulose membrane. Replicate membranes were probed with anti-phospho-acetyl CoA carboxylase (pACC; S79), anti-GFP or anti-SnRK1α1 antibodies. Protein bands were quantified using the Image Studio Live 5.2 software (LI-CORE). The signal ratio pACC/GFP was determined to normalize for differences in loading and construct expression and used as readout of SnRK1 activity. A 4-h extended night (ZT28) sample was loaded in every gel to normalize for technical variance. Loaded samples: 1-3: ZT8, 4-6: ZT16 (night), 7-9: ZT28 (extended night). Protein loading was visualized by staining with Ponceau Red (the gel number indicates which set of Ponceau Red stained images corresponds to which SnRK1 reporter construct and antibody).

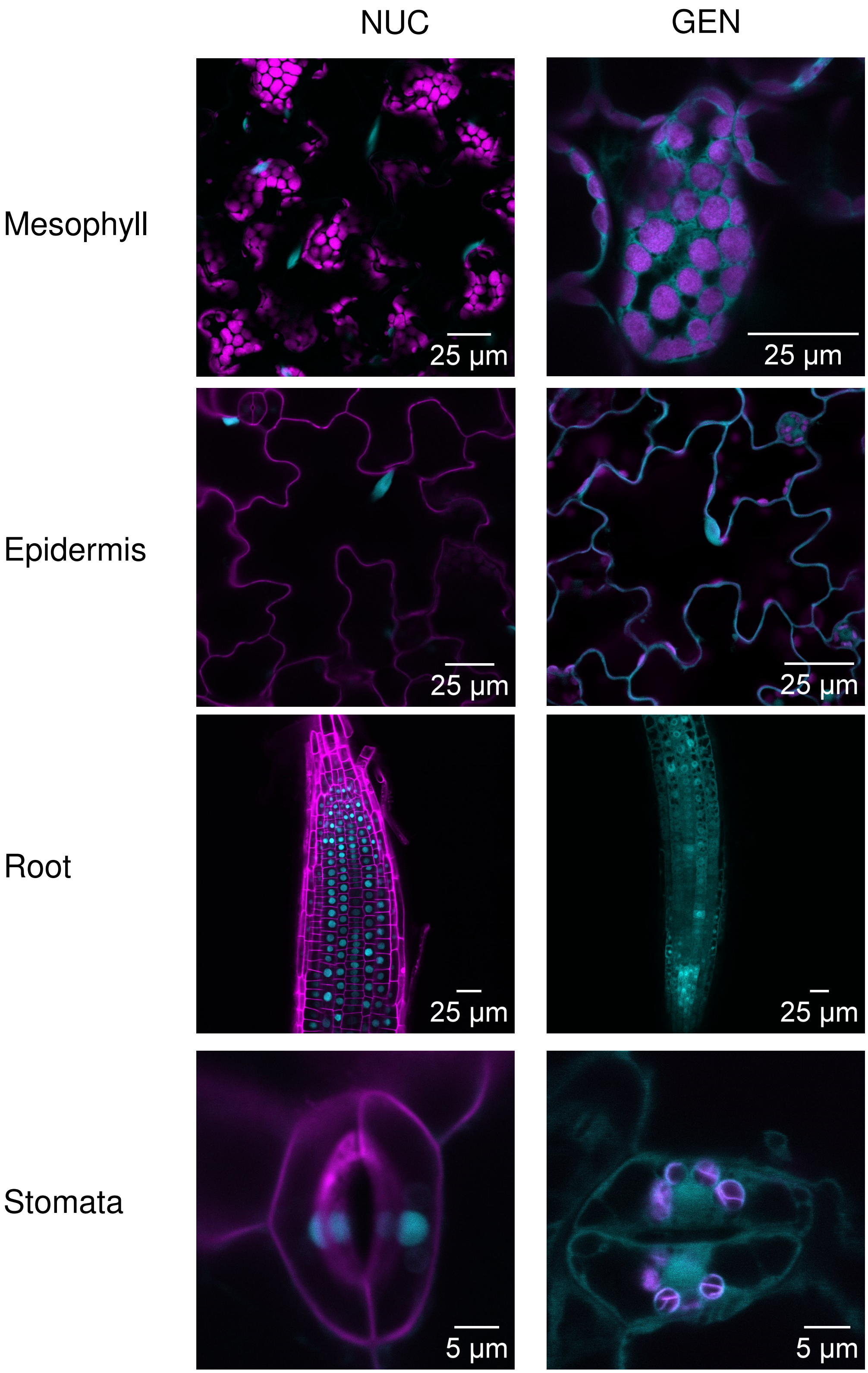


**Supplementary Figure S2: Localization of reporter peptides by confocal microscopy.**

Seedlings of NUC and GEN lines were grown on MS plates for 6 days and GFP signal was visualized by confocal microscopy, as described in Methods. NUC seedlings were further treated with propidium iodide (PI) to stain the cell wall. GEN samples were not stained with PI, to avoid overlapping signals between PI and plastids (chlorophyll). The figures show merged images of GFP and chlorophyll autofluorescence from the indicated tissues.

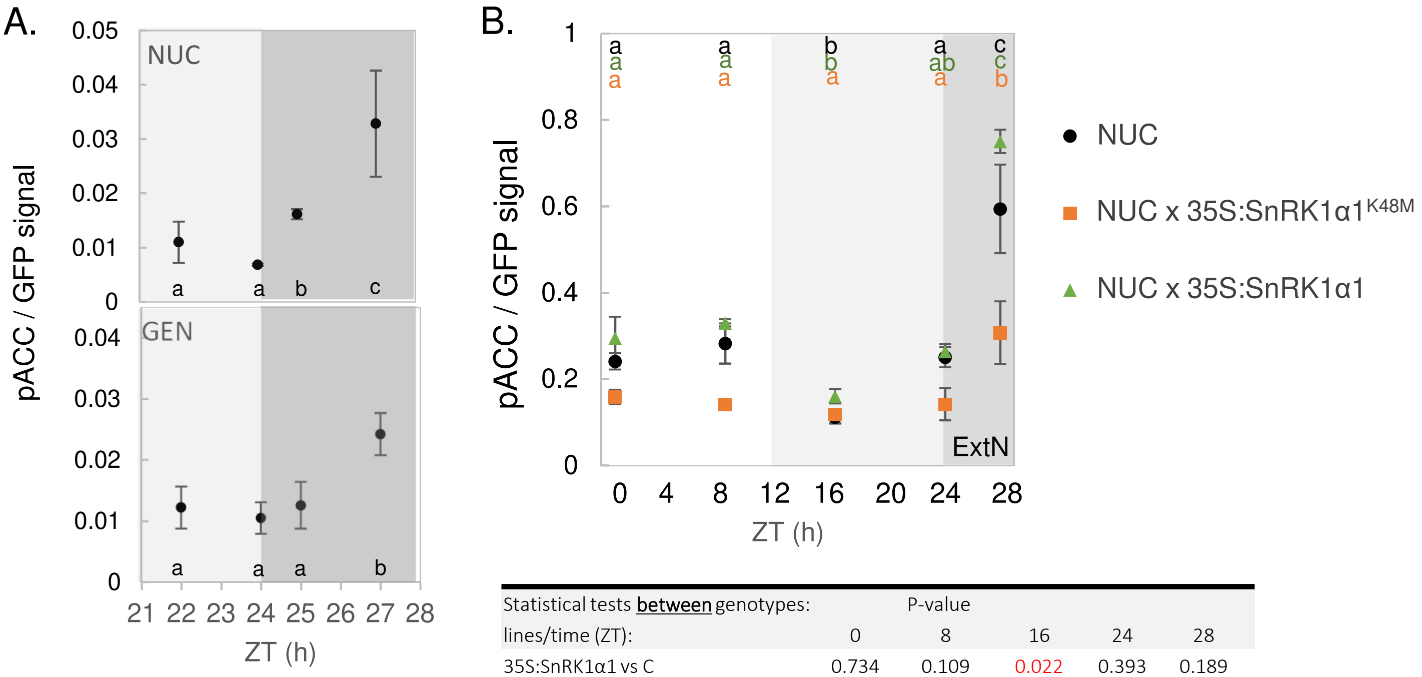


| Statistical tests **between** genotypes: | P-value | | | | |
|---|---|---|---|---|---|
| lines/time (ZT): | 0 | 8 | 16 | 24 | 28 |
| 35S:SnRK1α1 vs C | 0.734 | 0.109 | 0.022 | 0.393 | 0.189 |
| 35S:SnRK1α1$^{K48M}$ vs C | 0.023 | 0.001 | 0.892 | 0.003 | 0.006 |
| 35S:SnRK1α1 vs 35S:SnRK1α1$^{K48M}$ | 0.002 | 0.001 | 0.004 | 0.001 | <0.001 |

C.

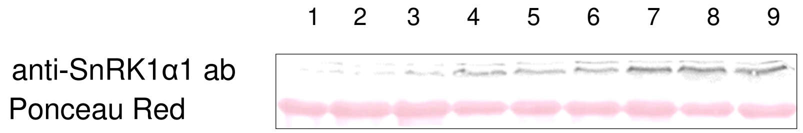


**Supplementary Figure S3: Validation of the NUC and GEN constructs in whole Arabidopsis rosettes**

(A) Extended darkness. NUC and GEN lines were grown in long day (16-h light / 8-h dark) conditions, with 160 µmol m$^{-2}$ s$^{-1}$ irradiance, for 22 days after sowing (22 DAS). On the day of harvest, plants were subjected to 3 h of extended night to activate SnRK1, and sampled at: ZT 20, 22, 24, 25, and 27. Immunoblots were performed as described in Methods and in Supplementary Figure S1C. Top -NUC, bottom – GEN. Letters represent statistical analysis (one-way ANOVA, followed by pairwise multiple comparison post-testing using the Holm-Sidak method, *P*<0.05). Results are shown as mean ± SD (n=2-4 biological replicates). See also Supplementary Dataset S1 – Exp.1. (B-C) High and low activity SnRK1 mutants. (B) NUC, NUC x 35S:SnRK1$^{K48M}$ and NUC x 35S:SnRK1α1 (i.e., overexpressing and dominant negative SnRK1 mutant lines crossed to NUC) were grown in equinoctial growth conditions (12-h light / 12-h dark) with 160 µmol m$^{-2}$ s$^{-1}$ irradiance for 22 DAS. Plants were harvested throughout day 23 after sowing at 8-h intervals and after 4 h of extended night (ExtN). Four replicates (each of 3-5 plants) were harvested at each time point. Immunoblots were performed as described in Methods and in Supplementary Figure S1C. Results are shown as mean ± SD (n=4 biological replicates). Statistical analysis: significance was tested separately for each line using one-way ANOVA, followed by pairwise multiple comparison post-testing using the Holm-Sidak method (*P* < 0.05). Letters represent significant changes between different times for a given genotype. The statistical analysis for comparison between lines at a given time point is presented in the table, with significant (*P*<0.05) differences highlighted in red. See also Supplementary Dataset S1 – Exp.2. (C) Immunoblot assay using anti-SnRK1α1 antibody to confirm expression of the introduced SnRK1α1 and SnRK1$^{K48M}$ proteins in the overexpression lines. All samples were harvested from the respective lines at ZT28 (4-h extended night): 1-3, NUC; 4-6, NUC x 35S:SnRK1α1$^{K48M}$; 7-9 NUC x 35S:SnRK1α1.

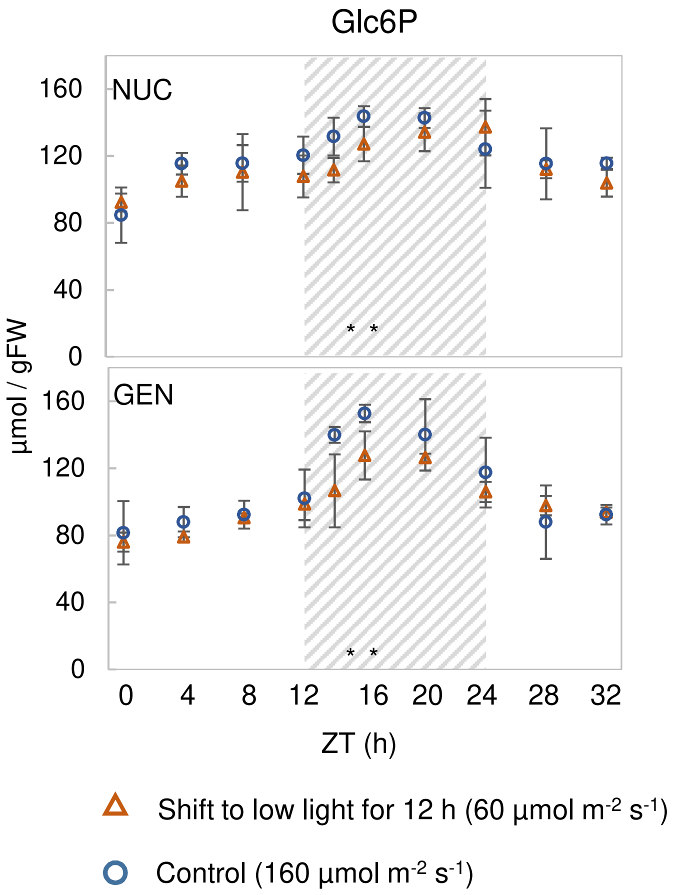


**Supplementary Figure S4: Comparison of different low light treatments (supplementary to Figures 2 and 3).**

Rosette Glc6P levels in NUC (upper panel) and GEN (lower panel) following transfer to continuous low light (90 µmol $m^{-2}$ $s^{-1}$ ), compared to a control in growth conditions (12-h photoperiod, 160 µmol $m^{-2}$ $s^{-1}$ irradiance) (n.b. data are from the same experiment shown in Figure 2). Results are shown as mean ± SD (n=3-4 biological replicates). Statistical analysis: one-way ANOVA, followed by pairwise multiple comparison post-testing using the Holm-Sidak method. Asterisks represent significant differences ($P$<0.05) between lines at a given time point. See also Supplementary Dataset S1 – Exp.3-Exp.4.

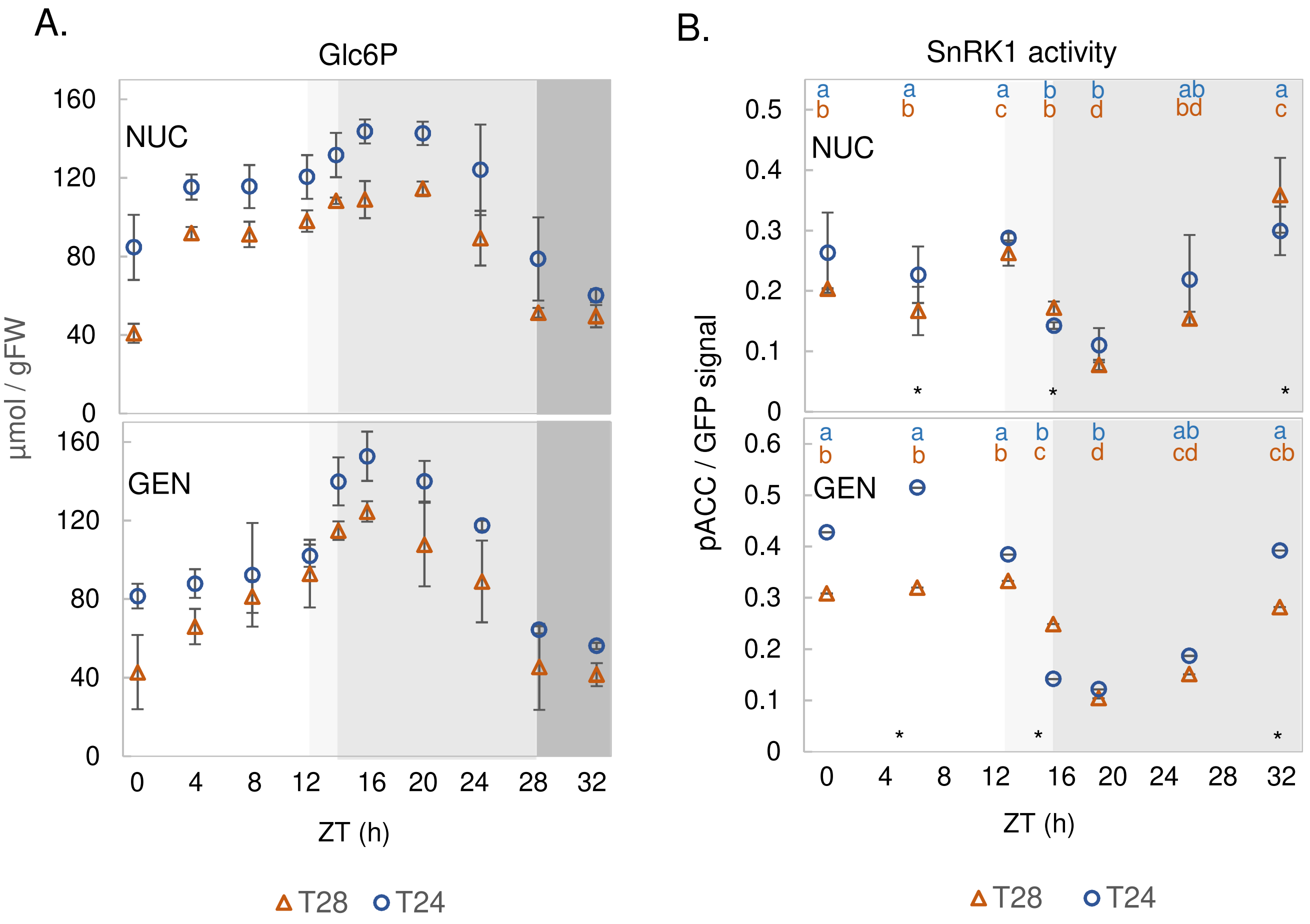


**Supplementary Figure S5: T28 and T24 cycles; diel Glc6P levels and comparison of SnRK1 activity excluding time points affected by C starvation (supplementary to Figure 4).**

(A) Rosette Glc6P levels from the T28 cycle experiment (described in Figure 4; Supplementary Dataset S1 Exp.5) for the NUC (upper) and GEN (lower) reporter lines. (B) Focused comparison (ZT4-ZT24) of SnRK1 activity between T28 and the T24 control, excluding ZT0, ZT28 and ZT32 when plants were C-starved in the T28 cycle. Upper panel NUC line, lower panel, GEN line. In (A) and (B), results are shown as mean ± SD (n=3 biological replicates), and the white background represents times when plants were in the light in both T24 control and T28 cycles, pale grey denotes darkness in the T24 control and light in the T28 cycle, mid-grey denotes darkness in both T-cycles, and dark grey represents extended night in both T cycles. Statistical analysis was done by three separate one-way ANOVA tests, followed by pairwise multiple comparison post-testing using the Holm-Sidak method ($P< 0.05$): (i) comparing changes with time within the control (T24) line, indicated by blue letters; (ii) comparing changes with time within the T28 line, indicated by orange letters; and (iii) comparing between T24 and T28 samples at a given time point, indicated by asterisks.

A.

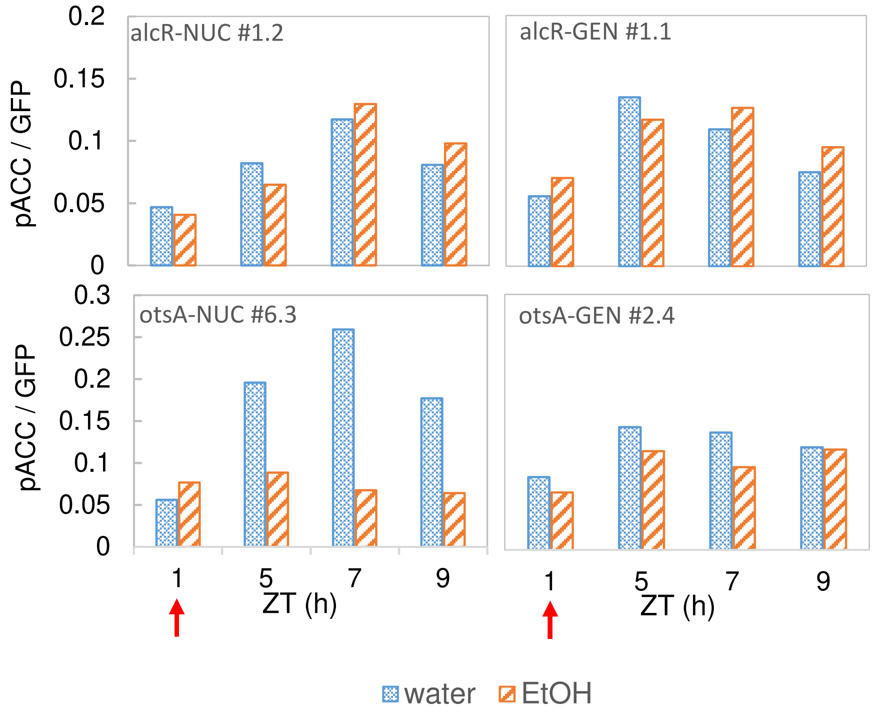


B.

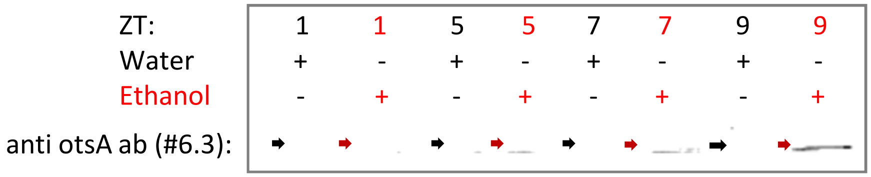


**Supplementary Figure S6: Response of NUC and GEN phosphorylation in crosses with a line containing inducible bacterial TPS**.

NUC and GEN reporter lines were crossed with an ethanol-inducible otsA (bacterial TPS) line and with the empty vector (alcR) control line, as described in Methods. Selected homozygous plants (#6.3 NUC x iotsA; #2.4 GEN x iotsA, alcR #1.1-1.2) were grown in long day conditions (16-h light / 8 h dark) under 160 µmol $m^{-2}$ $s^{-1}$ illumination for 20 days. On day 21 after sowing, plants were sprayed at ZT1 (red arrow) with either water (mock induction control) or with 2% (v/v) ethanol (EtOH) to induce the expression of the otsA protein and transiently increase Tre6P levels. Two plants (whole rosettes) were harvested and pooled as representative samples at 4, 6 and 8-h after water/EtOH spraying. (A) changes in SnRK1 phosphorylation activity upon otsA (TPS) induction. The ratios between the ethanol-induced and water control were determined for each time point, and then combined across all time points to test for significance in a given line. The differences were significant (Student's t-test, $P<0.05$) for #6.3 NUC x iotsA but not for #2.4 GEN x iots or the control alcR lines. (B) Immunoblot to confirm the inducible accumulation of otsA after spraying with ethanol in line #6.3 NUC x iotsA (indicated by the red arrows).

**Supplementary Figure S7: Impact of transiently elevated Tre6P in the light period on the levels of selected metabolites (supplementary to Figure 5).**

Line #6.3 (NUC x iotsA) was grown in equinoctial growth conditions (12-h light / 12 h dark) with 160 µmol $m^{-2}$ $s^{-1}$ irradiance for 20 days. On day 21 after sowing, plants were sprayed with either water or 2% (v/v) ethanol to induce the expression of bacterial TPS (otsA) at ZT4 (red arrow), and then harvested at ZT7, ZT9 and ZT11. Depicted are changes in selected metabolites. 2-OG, 2-oxoglutarate. Results are shown as mean ± SD (n = 3-4 biological replicates). Asterisks indicate statistically significant (*P*<0.05) differences between the water and ethanol-treated samples by one-way ANOVA followed by pairwise multiple comparison post-testing using the Holm-Sidak method. Data for additional metabolites from the same experiment are shown in Supplementary Dataset S1 Exp.7.

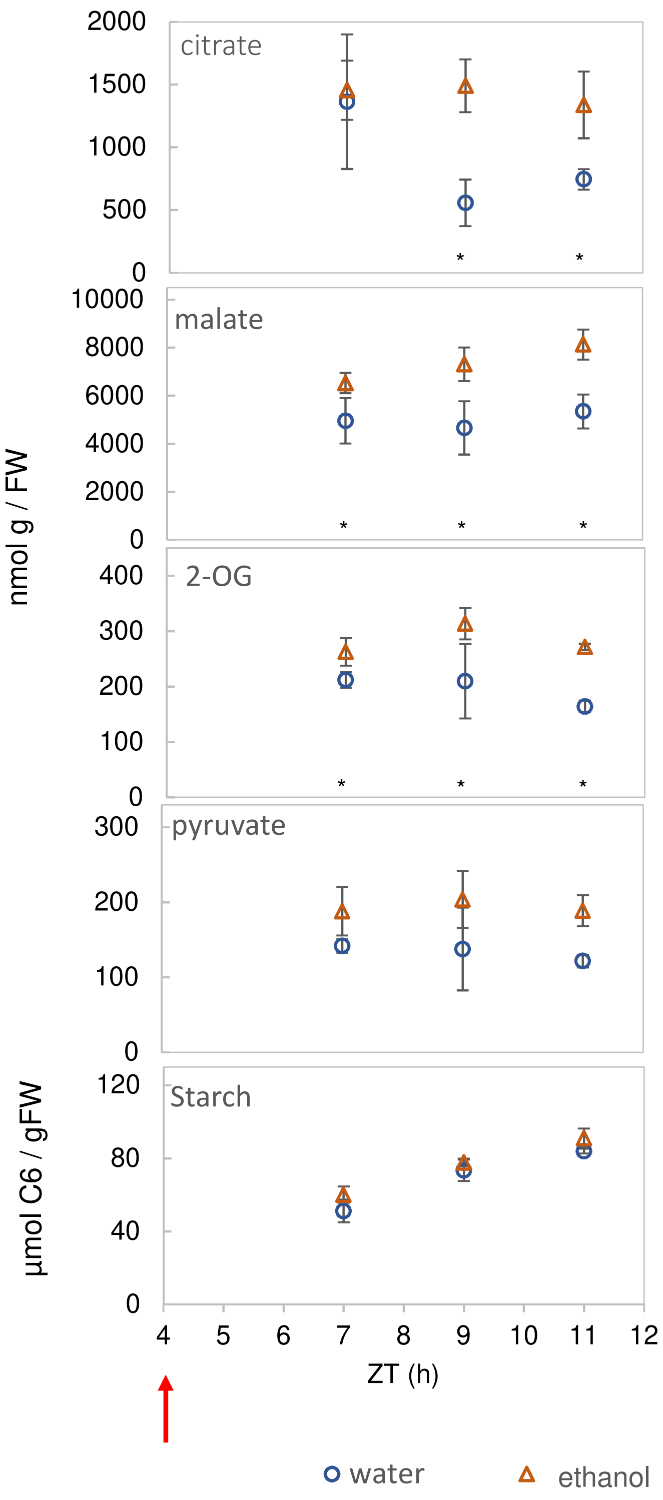

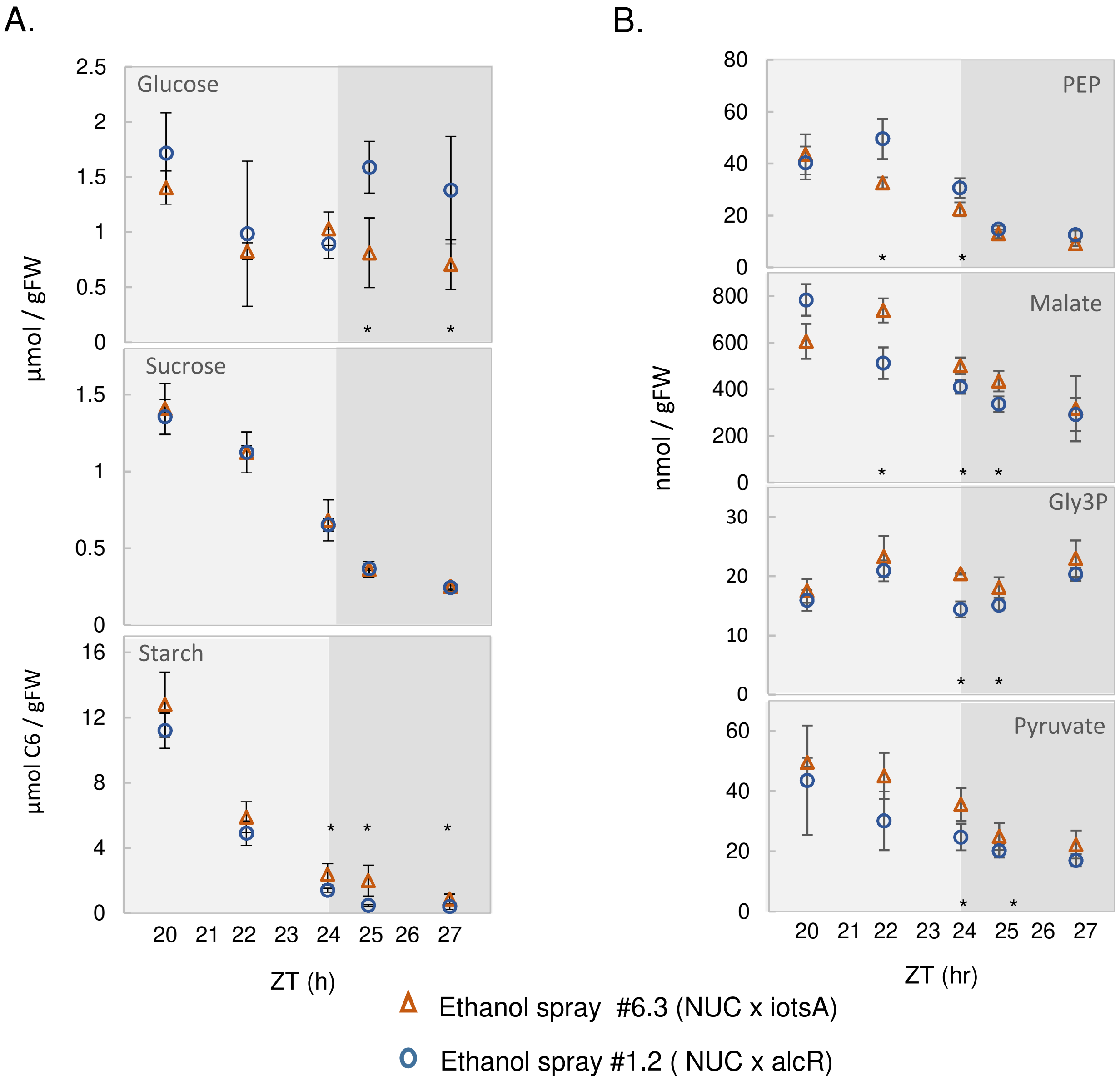


**Supplementary Figure S8: Transient elevation of Tre6P at the end of the night slows down starch mobilization and leads to low glucose and high organic acid levels (supplementary to Figure 6).**

Line #6.3 (NUC x iotsA) and control line #1.2 (NUC x alcR) were grown in short day conditions (6-h light / 18-h dark) with 160 µmol $m^{-2}$ $s^{-1}$ irradiance for 24 days. On day 25 after sowing, plants were sprayed at ZT20 with 2% (v/v) ethanol to induce expression of the otsA (TPS) protein, and thereby transiently increase Tre6P levels. After samples were harvested towards EN at ZT22 and ZT24, the remaining plants were transferred to extended darkness and harvested at ZT25 and ZT27 (as described in Figure 6). Four plants (whole rosettes) were pooled for each biological replicate. Rosettes were extracted for measurements of: (A) soluble sugars and starch, and (B) PEP, malate, glycerol 3-phosphate (Gly3P) and pyruvate. Data for additional metabolites are shown in Supplementary Figure S8, and Supplementary Dataset S1 – Exp.7. Results are shown as mean ± SD (n = 3-4 biological replicates). Asterisks indicate significant (P<0.05) differences between the #6.3 (NUC x iotsA) and control lines according to one-way ANOVA with post-testing using the Holm-Sidak method.

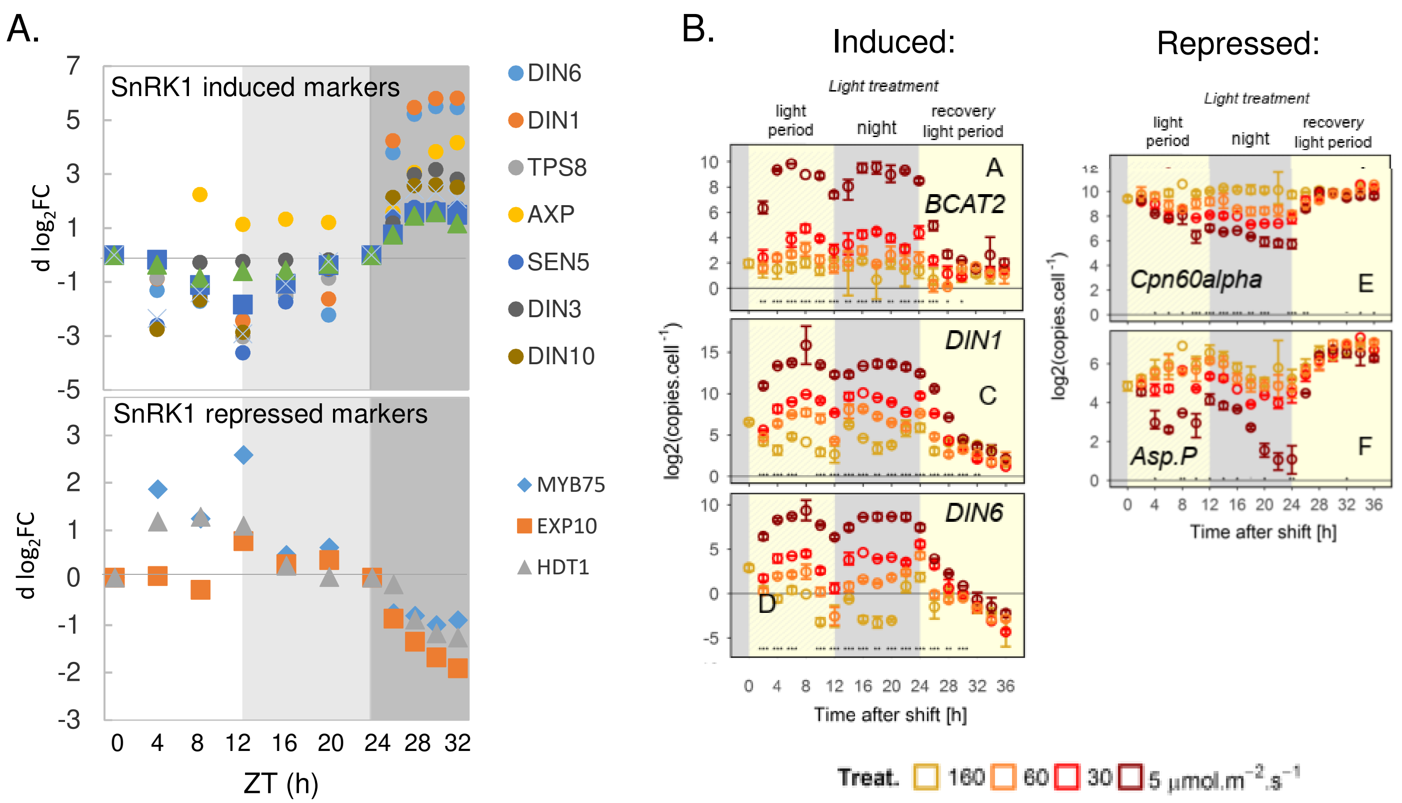


**Supplementary Figure S9: Transcript abundance for SnRK1 marker genes during a diel cycle, after an extension of the night and after a sudden decrease in light intensity for one day.**

Publicly available gene expression datasets were mined for selected genes that are commonly referred to as SnRK1 marker genes, based on their response to transient over-expression of SnRK1α1 in Arabidopsis mesophyll protoplasts (Baena-González *et al.*, 2007). (A). Transcript abundance in wild-type Columbia-0 rosettes during a diel cycle in equinoctial conditions at an irradiance of 160 µmol $m^{-2}$ $s^{-1}$ and after an extension of the night. Expression data were extracted from the ATH1 microarray datasets reported in Usadel et al. (2008) and were normalized to the respective expression level at ZT0. (B) Transcript abundance in response to sudden low light day. Grey – night, dark grey – extended night. The experiment was conducted essentially as in Figure 3, except that a wider range of low light intensities was used. Wild-type Columbia-0 plants were grown in equinoctial growth conditions (12-h photoperiod) with 160 µmol $m^{-2}$ $s^{-1}$ irradiance for 19 days. At the onset of the light period on day 20 after sowing, irradiance was reduced to 60, 30 or 5 µmol $m^{-2}$ $s^{-1}$ for the 12-h light period (low-light day). The plants were darkened at ZT12, and re-illuminated at ZT24 at the original growth irradiance of 160 µmol $m^{-2}$ $s^{-1}$ (recovery day). Results are shown as mean ± 95% confidence limits (n = 3 biological replicate, each containing five pooled rosettes). The symbols just above the x-axis indicated statistical significance of the combined response according to ANOVA: *** $P$=0, ** $P$<0.001, * P<0.01, · $P$<0.05. $P$-values are adjusted using Benjamini & Hochberg false discovery rate across all time points in a given trait; dashes (-) indicate when significant differences were rejected by the Tukey's honest significant difference post-test. Data were extracted from Supplementary Figure S6 in Moraes *et al.* (2019).
