## Supplementary Tables S1-S6 for "Diel fluctuations in *in-vivo* SnRK1 activity in Arabidopsis rosettes during light-dark cycles"

**Supplementary Table S1:** Correlations between metabolite levels and NUC or GEN phosphorylation, performed separately for the light period and the night (supplementary to Table 1 and Figure 1).

$R^2$ and *P* values are shown for linear regression analyses performed separately on data from the three independent control experiments described in Figure 1. Regressions were performed using data for individual replicates from the light period (ZT4, ZT8, ZT12) or the night (ZT16, ZT20, ZT24). Linear regression analysis was performed using the least squares method. The column +/- indicates whether the slope of the regression is positive (+) or negative (-). Significant correlations are highlight in red (*P*<0.05). Individual plots are provided in Supplementary Dataset S2.

| Experiment | Reporter | Metabolite | Day | | | Night | | |
|---|---|---|---|---|---|---|---|---|
| | | | $R^2$ | *P*-value | +/- | $R^2$ | *P*-value | +/- |
| Exp. 1 | NUC | Tre6P | 0.010 | 0.232 | + | 0.108 | 0.232 | - |
| | NUC | Glc6P | 0.035 | 0.558 | - | 0.595 | $2.7\times10^{-4}$ | - |
| | NUC | Glc1P | 0.126 | 0.257 | - | 0.609 | $6.0\times10^{-4}$ | - |
| | NUC | Starch | 0.000 | 1.000 | - | 0.822 | $3.2\times10^{-6}$ | - |
| | NUC | Sucrose | 0.000 | 0.973 | + | 0.760 | $2.3\times10^{-5}$ | - |
| | NUC | Glucose | 0.004 | 0.854 | + | 0.280 | 0.043 | - |
| | GEN | Tre6P | 0.240 | 0.181 | - | 0.347 | 0.044 | - |
| | GEN | Glc6P | 0.184 | 0.249 | - | 0.478 | 0.013 | - |
| | GEN | Glc1P | 0.099 | 0.410 | - | 0.243 | 0.103 | - |
| | GEN | Starch | 0.139 | 0.322 | - | 0.547 | 0.006 | - |
| | GEN | Sucrose | 0.241 | 0.180 | - | 0.348 | 0.004 | - |
| | GEN | Glucose | 0.172 | 0.267 | + | 0.275 | 0.080 | + |
| Exp. 2 | NUC | Tre6P | 0.152 | 0.300 | + | 0.441 | 0.007 | - |
| | NUC | Glc6P | 0.053 | 0.550 | - | 0.114 | 0.219 | - |
| | NUC | Glc1P | 0.251 | 0.169 | - | 0.364 | 0.017 | - |
| | NUC | Starch | 0.052 | 0.554 | + | 0.644 | 0.001 | - |
| | NUC | Sucrose | 0.124 | 0.352 | + | 0.514 | 0.006 | - |
| | NUC | Glucose | 0.082 | 0.454 | + | 0.006 | 0.808 | - |
| | GEN | Tre6P | 0.154 | 0.108 | - | 0.656 | $4.5\times10^{-4}$ | - |
| | GEN | Glc6P | 0.112 | 0.378 | - | 0.536 | 0.003 | - |
| | GEN | Glc1P | 0.122 | 0.356 | - | 0.707 | $1.7\times10^{-4}$ | - |
| | GEN | Starch | 0.003 | 0.892 | - | 0.740 | $7.9\times10^{-5}$ | - |
| | GEN | Sucrose | 0.249 | 0.171 | - | 0.824 | $7.4\times10^{-6}$ | - |
| | GEN | Glucose | 0.461 | 0.044 | + | 0.148 | 0.175 | - |
| Exp. 3 | NUC | Tre6P | 0.022 | 0.644 | - | 0.278 | 0.066 | - |
| | NUC | Glc6P | 0.022 | 0.811 | + | 0.251 | 0.057 | - |
| | NUC | Glc1P | 0.012 | 0.737 | + | 0.078 | 0.113 | - |
| | NUC | Starch | 0.001 | 0.969 | + | 0.686 | $7.3\times10^{-5}$ | - |
| | NUC | Sucrose | 0.056 | 0.458 | + | 0.011 | 0.702 | - |
| | NUC | Glucose | 0.001 | 0.917 | - | 0.131 | 0.168 | - |

**Supplementary Table S2**: Correlations between metabolite levels (supplementary to Table 1 and Figure 1).

$R^2$ and *P*-values are shown for linear regression analyses performed separately on data from the three independent control experiments with the NUC line and two independent experiments with the GEN line described in Figure 1 (12-h photoperiod, 160 µmol $m^{-2}$ $s^{-1}$ irradiance). Regressions were performed using data for individual replicates from the complete diel cycle (ZT0 to ZT24). The linear regression analysis was done by using the least squares method. The column +/- indicates whether the slope of the regression is positive (+) or negative (-). Significant correlations are highlight in red (*P*<0.05), and metabolite pairs highlighted in bold were significantly correlated in all three NUC experiments and in the two independent GEN experiments (not presented). Individual plots are provided in Supplementary Dataset S2.

| **NUC** | | **Experiment 1** | | | **Experiment 2** | | | **Experiment 3** | | |
|---|---|---|---|---|---|---|---|---|---|---|
| | | **$R^2$** | ***P*-value** | **+/-** | **$R^2$** | ***P*-value** | **+/-** | **$R^2$** | ***P*-value** | **+/-** |
| Tre6P | Glc6P | 0.532 | $1.1x10^{-5}$ | + | $1x10^{-4}$ | 0.885 | - | 0.287 | 0.006 | + |
| Tre6P | Glc1P | 0.340 | 0.001 | + | 0.007 | 0.697 | + | 0.147 | 0.048 | + |
| **Tre6P** | **Sucrose** | 0.400 | $3.1x10^{-4}$ | + | 0.821 | $2.7\ x10^{-9}$ | + | 0.247 | 0.010 | + |
| **Tre6P** | **Starch** | 0.372 | 0.001 | + | 0.676 | $1.5\ x10^{-6}$ | + | 0.432 | $2.7\ x10^{-4}$ | + |
| **Glc1P** | **Glc6P** | 0.860 | $1.3x10^{-12}$ | + | 0.654 | $2.2\ x10^{-6}$ | + | 0.279 | 0.007 | + |
| Glc6P | Glucose | 0.011 | 0.604 | + | 0.011 | 0.637 | - | 0.048 | 0.284 | + |
| Glc6P | Sucrose | 0.645 | $2.7x10^{-7}$ | + | $4x10^{-4}$ | 0.924 | + | 0.529 | $3.8\ x10^{-5}$ | + |
| Glc6P | Starch | 0.586 | $2.1x10^{-6}$ | + | 0.001 | 0.885 | + | 0.695 | $2.3\ x10^{-7}$ | + |
| Glc1P | Starch | 0.601 | $1.3\ x10^{-6}$ | + | 0.051 | 0.300 | + | 0.089 | 0.138 | + |
| **Sucrose** | **Starch** | 0.642 | $3.0\ x10^{-7}$ | + | 0.775 | $3.1\ x10^{-8}$ | + | 0.474 | $1.0\ x10^{-4}$ | + |
| Glucose | Starch | 0.011 | 0.537 | + | 0.094 | 0.156 | + | 0.019 | 0.491 | + |

| **GEN** | | **Experiment 1** | | | **Experiment 2** | | |
|---|---|---|---|---|---|---|---|
| | | **$R^2$** | ***P*-value** | **+/-** | **$R^2$** | ***P*-value** | **+/-** |
| Tre6P | Glc6P | 0.532 | $1.1\ x10^{-5}$ | + | $1x10^{-4}$ | 0.885 | - |
| Tre6P | Glc1P | 0.340 | 0.001 | + | 0.007 | 0.697 | + |
| Tre6P | Sucrose | 0.400 | $3.1\ x10^{-4}$ | + | 0.821 | $2.7\ x10^{-9}$ | + |
| Tre6P | Starch | 0.372 | 0.001 | + | 0.676 | $1.5\ x10^{-6}$ | + |
| Glc1P | Glc6P | 0.860 | $1.3\ x10^{-12}$ | + | 0.654 | $2.2\ x10^{-6}$ | + |
| Glc6P | Glucose | 0.011 | 0.604 | + | 0.011 | 0.637 | - |
| Glc6P | Sucrose | 0.645 | $2.7\ x10^{-7}$ | + | $4x10^{-4}$ | 0.924 | + |
| Glc6P | Starch | 0.586 | $2.1\ x10^{-6}$ | + | 0.001 | 0.885 | + |
| Glc1P | Starch | 0.601 | $1.3\ x10^{-6}$ | + | 0.051 | 0.300 | + |
| Sucrose | Starch | 0.642 | $3.0\ x10^{-7}$ | + | 0.775 | $3.1\ x10^{-8}$ | + |
| Glucose | Starch | 0.011 | 0.537 | + | 0.094 | 0.156 | + |

**Supplementary Table S3:** Correlations between changes in metabolite levels and changes in NUC or GEN phosphorylation (supplementary to Table 1 and Figure 1).

$R^2$ and *P*-values are shown for linear regression analyses performed separately on data from the three independent control experiments described in Figure 1. Changes in activity were calculated as the difference between absolute levels (averages) of two consecutive time points, divided by the time difference (h). The linear regression analysis was done by using the least squares method. Significant correlations are highlighted in red ($P<0.05$). The column +/- indicates whether the slope of the regression is positive (+) or negative (-).

| Reporter | Metabolite | $R^2$ | *P*-value | +/- |
|---|---|---|---|---|
| NUC | Tre6P | 0.003 | 0.845 | + |
| NUC | Glc6P | 0.252 | 0.029 | - |
| NUC | Glc1P | 0.399 | 0.004 | - |
| NUC | Starch | 0.037 | 0.429 | + |
| NUC | Sucrose | 0.163 | 0.087 | + |
| NUC | Glucose | 0.009 | 0.706 | - |
| GEN | Tre6P | 0.295 | 0.068 | - |
| GEN | Glc6P | 0.617 | 0.002 | - |
| GEN | Glc1P | 0.369 | 0.028 | - |
| GEN | Starch | 0.010 | 0.743 | - |
| GEN | Sucrose | 0.121 | 0.244 | - |
| GEN | Glucose | 0.019 | 0.657 | + |

**Supplementary Table S4:** Correlations between metabolite levels and NUC or GEN phosphorylation in growth regimes that differed from the equinoctial T24 cycle (supplementary to Figures 3 and 4).

Continuous (90) is the experiment reported in Figure 3 and Supplementary Dataset S1 Exp.4, in which plants were grown in a 12-h photoperiod at 160 µmol $m^{-2}$ $s^{-1}$ irradiance and then transferred at dawn to continuous irradiance at 90 µmol $m^{-2}$ $s^{-1}$ for 32 h. Regressions were performed using data from individual replicates. T28 denotes the experiment reported in Figure 4 and Supplementary Dataset S2 Exp.5, in which plants were grown in a 14-h light / 14-h dark cycle. Regression analyses was performed using data from individual replicates on all times in light dark cycle (i.e. from ZT0 to ZT28). T28 (ZT4-ZT24) shows an additional analysis from the T28 experiment, using only time points between ZT4-ZT24 to eliminate possible interference due to C starvation at ZT0 and ZT28 (see Figure 4). For data from both experiments, linear regression analysis was done using the least squares method. The column +/- indicates whether the slope of the regression is positive (+) or negative (-). Significant correlations are highlighted in red (*P*<0.05). Individual plots are provided in Supplementary Dataset S2. n.d., metabolite not determined in this experiment.

| Reporter | Metabolite | Continuous (90) | | | T28 | | | T28 (ZT4-ZT24) | | |
|---|---|---|---|---|---|---|---|---|---|---|
| | | $R^2$ | *P*-value | +/- | $R^2$ | *P*-value | +/- | $R^2$ | *P*-value | +/- |
| NUC | Tre6P | 0.462 | $2.6x10^{-4}$ | - | n.d. | n.d. | | n.d. | | |
| NUC | Glc6P | 0.414 | 0.001 | - | 0.889 | $1.9x10^{-13}$ | - | 0.366 | 0.004 | - |
| NUC | Glc1P | 0.228 | 0.018 | - | n.d. | n.d. | | n.d. | | |
| NUC | Starch | 0.012 | 0.972 | - | 0.534 | $1.5x10^{-5}$ | - | 0.255 | 0.020 | - |
| NUC | Sucrose | 0.057 | 0.243 | - | 0.687 | $9.4x10^{-8}$ | - | 0.151 | 0.082 | - |
| NUC | Glucose | 0.059 | 0.738 | - | 0.027 | 0.416 | | 0.002 | 0.857 | - |
| GEN | Tre6P | 0.337 | 0.004 | - | n.d. | n.d. | | n.d. | n.d. | |
| GEN | Glc6P | 0.314 | 0.005 | - | 0.674 | $2.8x10^{-7}$ | - | 0.496 | 0.001 | - |
| GEN | Glc1P | 0.262 | 0.013 | - | n.d. | n.d. | | n.d. | n.d. | |
| GEN | Starch | 0.083 | 0.322 | - | 0.477 | $6.8x10^{-5}$ | - | 0.019 | 0.550 | - |
| GEN | Sucrose | 0.021 | 0.511 | - | 0.693 | $7.2x10^{-5}$ | - | 0.065 | 0.265 | - |
| GEN | Glucose | 0.097 | 0.099 | - | 0.260 | 0.007 | - | 0.115 | 0.132 | + |

**Supplementary Table S5**: Summary of correlation analysis between changes in SnRK1 NUC and GEN activity to changes in metabolite levels (supplementary to Figures 3 and 4)

See Supplementary Table S3 for details of the experiments and the time spans used for the correlation analysis. Changes in activity were calculated as the difference between absolute levels (averages) of two consecutive time points, divided by the time difference (h). The linear regression analysis was done using the least squares method. Significant correlations ($P<0.05$) are highlighted in red. The column +/- indicates whether the slope of the regression is positive (+) or negative (-). n.d., metabolite not determined in this experiment.

| Reporter | Metabolite | Continuous (90) | | | T28 | | | T28 (ZT4-ZT24) | | |
|---|---|---|---|---|---|---|---|---|---|---|
| | | $R^2$ | *P*-value | +/- | $R^2$ | *P*-value | +/- | $R^2$ | *P*-value | +/- |
| NUC | Tre6P | 0.542 | 0.059 | - | n.d. | n.d. | | n.d. | n.d. | |
| NUC | Glc6P | 0.550 | 0.056 | - | 0.873 | 0.001 | - | 0.444 | 0.148 | - |
| NUC | Glc1P | 0.390 | 0.134 | - | n.d. | n.d. | | n.d. | n.d. | |
| NUC | Starch | 0.114 | 0.459 | + | 0.191 | 0.279 | - | 0.051 | 0.668 | - |
| NUC | Sucrose | 0.044 | 0.650 | + | 0.320 | 0.144 | - | 0.005 | 0.899 | + |
| NUC | Glucose | 0.131 | 0.426 | - | 0.015 | 0.771 | - | 0.026 | 0.758 | + |
| GEN | Tre6P | 0.403 | 0.125 | - | n.d. | n.d. | | n.d. | n.d. | |
| GEN | Glc6P | 0.460 | 0.094 | - | 0.537 | 0.039 | - | 0.485 | 0.124 | - |
| GEN | Glc1P | 0.422 | 0.114 | - | n.d. | n.d. | | n.d. | n.d. | |
| GEN | Starch | 0.304 | 0.199 | - | 0.178 | 0.298 | - | 0.030 | 0.744 | - |
| GEN | Sucrose | 0.097 | 0.496 | - | 0.391 | 0.097 | - | 0.001 | 0.950 | + |
| GEN | Glucose | 0.204 | 0.309 | - | 0.334 | 0.134 | - | 0.074 | 0.602 | + |

**Supplementary Table S6**: Correlation analysis between metabolite levels and NUC phosphorylation following transient elevation of Tre6P in the light period (supplementary to Figure 5, Supplementary Dataset S2 Exp.7).

$R^2$ and *P*-values are shown for regression analyses after treatment with ethanol, or in the water (mock induction) control. Linear regression analyses were performed using the least squares method on data for individual replicates. The column +/- indicates whether the slope of the regression is positive (+) or negative (-). Significant correlations (*P*<0.05) are highlighted in red.

| **Reporter** | **Metabolite** | **Water** | | **Ethanol** | |
|---|---|---|---|---|---|
| | | **$R^2$** | ***P*-value** | **$R^2$** | ***P*-value** |
| NUC | Tre6P | 0.671 | 0.004 | 0.012 | 0.763 |
| NUC | Glc6P | 0.243 | 0.148 | 0.140 | 0.287 |
| NUC | Glc1P | 0.153 | 0.263 | 0.047 | 0.547 |
| NUC | Starch | 0.004 | 0.869 | 0.091 | 0.398 |
| NUC | Sucrose | 0.028 | 0.819 | 0.105 | 0.361 |
| NUC | Glucose | 0.000 | 0.983 | 0.328 | 0.083 |
