## Supplementary Text for "Diel fluctuations in *in-vivo* SnRK1 activity in Arabidopsis rosettes during light-dark cycles"

**Supplementary Text. Correlations between *in-vivo* SnRK1 activity and metabolites in continuous light, in T28 cycles and in experiments where TPS is induced.**

*In-vivo* SnRK1 activity correlates with Tre6P and especially Glc6P and Glc1P in a recurring equinoctial diel cycle (see main text, Table 1 and Supplementary Table S1). To test whether these correlations are found under perturbations of the diel cycle, we utilized the data from experiments where plants were shifted to continuous irradiance at 90 µmol $m^{-2}$ $s^{-1}$ µmol $m^{-2}$ $s^{-1}$ (Figure 3, Supplementary Dataset S1 Exp.4) or grown in a T28 cycle (Figure 4, Supplementary Dataset S1 Exp.5), and performed additional regressions between diel SnRK1 NUC and GEN phosphorylation and metabolite levels (Supplementary Table S3; see Supplementary Dataset S2 file for plots).

In the shift to constant lower irradiance, Tre6P, Glc6P and Glc1P were significantly and negatively correlated to SnRK1 NUC ($R^2$ = of 0.462, 0.414, and 0.2327, respectively) and GEN ($R^2$ of 0.3437, 0.314, and 0.262, respectively). No significant correlation was observed with starch, sucrose and glucose. Additional regressions were performed using derivatives (average change over time). These showed no significant correlation with any metabolite (Supplementary Table S4).

For plants growing in a T28 cycle (Fig. 4) the analysis was more restricted. Metabolites were only measured enzymatically and not by LC-MS/MS, so Tre6P and Glc1P were absent from these analyses. To avoid regressions that were driven by the very high phosphorylation of NUC and GEN at ZT0 and ZT28 (low C; Figure 4), the analyses were performed on two different sets of values; once over the full 28-h cycle and the second over a narrowed time scale from ZT4-ZT24. Regressions of absolute diel levels over the full 28-h cycle revealed significant correlation between NUC phosphorylation and Glc6P, starch and sucrose and between GEN phosphorylation and Glc6P, starch, sucrose and glucose. The more focused analysis using ZT4-ZT24 showed significant correlation between NUC phosphorylation and Glc6P and starch, and between GEN phosphorylation and Glc6P. Similarly, regressions of average change over time (derivatives) revealed a significant correlation between both NUC and GEN phosphorylation and Glc6P, using the full 28-h cycle (Supplementary Table S4).

Regressions were also calculated for the experiments with lines containing iTPS. These calculations were limited by the relatively small number of time points. When Tre6P was increased by TPS induction in the light period, (Fig. 5), NUC phosphorylation correlated negatively with Tre6P in the control ($R^2$ = 0.67, p = 0.76) but not in the induced TPS line ($R^2$ = 0.12, p = 0.004) (Supplementary Table S6). This is consistent with the idea that SnRK1 activity is regulated by a network including Tre6P, and that this network responds to decrease the contribution of Tre6P when Tre6P levels are artificially elevated. Glc was unrelated to SnRK1 in the control and negatively related after ethanol spraying ($R^2$ = 0 and 0.33, respectively), but was still only very weakly significant (*P* = 0.08). When Tre6P was elevated towards the end of the night and harvesting was continued into an extended night, NUC phosphorylation correlated negatively with Tre6P, Glc6P, Glc1P, sucrose and starch (not shown). This might reflect this experiment having been conducted at a time when C is becoming depleted and there is a decrease in the levels of many C metabolites that are potentially involved in regulation of SnRK1.
